# Sample-efficient time-resolved X-ray crystallography with serial small wedges

**DOI:** 10.64898/2026.10.04.756486

**Authors:** Sergei Bukhdruker, Roman Astashkin, Dmitry Bratanov, Nicolas Caramello, Sylvain Engilberge, Christian Baeken, Taras Balandin, Alexander Popov, Antoine Royant, Christoph Mueller-Dieckmann, Montserrat Soler-López, Ernst Bamberg, Valentin Gordeliy, Igor Melnikov

**Author notes:** Correspondence (S.B.); (V.G.); (I.M.).

## Abstract

Time-resolved crystallography can be inaccessible for membrane proteins and other difficult-to-crystallize targets for which suitable crystals are scarce. Here we present an acquisition strategy that records consecutive diffraction images during a small crystal rotation synchronized to a single activation pulse. Matching image segments are merged across crystals to reconstruct successive time windows, enabling a single crystal ensemble to capture an extended structural trajectory. Using the Mesh&Collect workflow at a synchrotron, we obtained ten consecutive 7.5-ms time windows at 2 Å resolution from 1,255 microcrystals of a bacteriorhodopsin triple mutant at room temperature. An additional 790 crystals extended the observation window to 500 ms. Rotation-based processing, retrospective time binning and dose control maximize information extracted from each crystal, while compatible measurements can be pooled across acquisitions. This approach makes millisecond time-resolved crystallography practical for crystal-limited targets at synchrotrons, with further development potentially extending the temporal resolution down to 1 ms.

## Introduction

Time-resolved crystallography connects molecular structure to function by measuring electron-density changes after reaction initiation.^1,2^ At synchrotrons, serial studies of membrane proteins have resolved proton transport in bacteriorhodopsin, inward proton pumping in xenorhodopsin, chloride transport, the prolonged photocycle of sensory rhodopsin II and channelrhodopsin activation, as well as ligand photopharmacology in a G protein-coupled receptor (GPCR).^3,4,5,6,7,8^ Complementary serial femtosecond crystallography (SFX) experiments at X-ray free-electron lasers (XFELs) have captured ultrafast retinal isomerization, sodium and chloride pumping, and early channelrhodopsin activation.^9,10,11,12,13,14,15^ These studies demonstrate the biological reach of serial crystallography, yet whether a time-resolved experiment is feasible can still depend on the availability of suitable crystals.

This constraint weighs more heavily on membrane proteins, for which expression, purification and crystallization may yield only a few well-diffracting crystals rather than a dense, reproducible suspension. For static structure determination, serial small-wedge crystallography already exploits such samples efficiently by combining rotation data from multiple crystals, as shown for GPCR structures.^16,17,18,19^ For time-resolved experiments, the challenge is to extract more temporal information from each usable crystal without exceeding its tolerable X-ray dose.

Aumonier *et al.* introduced time-resolved serial oscillation crystallography (TR-SOX), which records rotation series during continuous illumination of a blue-light photoreceptor and merges matching segments across 88 crystals into 63-ms time bins.^20^ Here, we combine this rotation-and-segmentation principle with the repeated-frame synchrotron sampling exemplified by Weinert *et al*.^3^, using single-pulse activation and short, dose-limited rotation wedges. Because continuous rotation data collection is synchronized with a short laser pulse, each fine-sliced diffraction image is acquired over a successive time interval during or after the activation pulse. Corresponding temporal segments are then merged from crystals with different orientations to reconstruct consecutive time windows.

Our acquisition strategy distributes the dose over multiple images within a small angular range, so that crystal rotation simultaneously defines both the diffraction orientation and the delay after activation. Experimental parameters are refined from each complete wedge, while corresponding image segments from many crystals are combined to achieve high completeness and improve the signal-to-noise ratio. Thus, each crystal contributes to multiple time windows, and a single crystal ensemble yields an extended structural trajectory. We implemented this approach with the Mesh&Collect workflow at the European Synchrotron Radiation Facility (ESRF) MASSIF-3 beamline.^21,22^ Using the T17A/T24A/T47A triple mutant of *Halobacterium salinarum* bacteriorhodopsin (BR_MT_), we obtained ten consecutive 7.5-ms datasets at 2 Å resolution from 1,255 crystals. Spectroscopic and structural controls established the sensitivity and operating range of the approach.

## Results

### A small rotation wedge records a structural trajectory

Starting at the activation trigger, each crystal is rotated through a small angle while consecutive diffraction images are recorded (Fig. 1a). We refer to the resulting small rotational dataset as a wedge, where both reaction time and reciprocal space are sampled. A segment comprises the consecutive images selected from a wedge or a full rotation dataset to reconstruct a time window (Fig. 1b). Merging corresponding segments from differently oriented crystals yields a series of time-resolved datasets (Fig. 1c). Thus, each crystal contributes to multiple time windows rather than being assigned to a single time window. The intermediate states indicated at the bottom of Fig. 1c are those expected over successive time windows guided by spectroscopy. Supplementary Figs. 1 and 2 illustrate a visual analogy and compare this sampling principle with other acquisition schemes.

**Figure 1.**
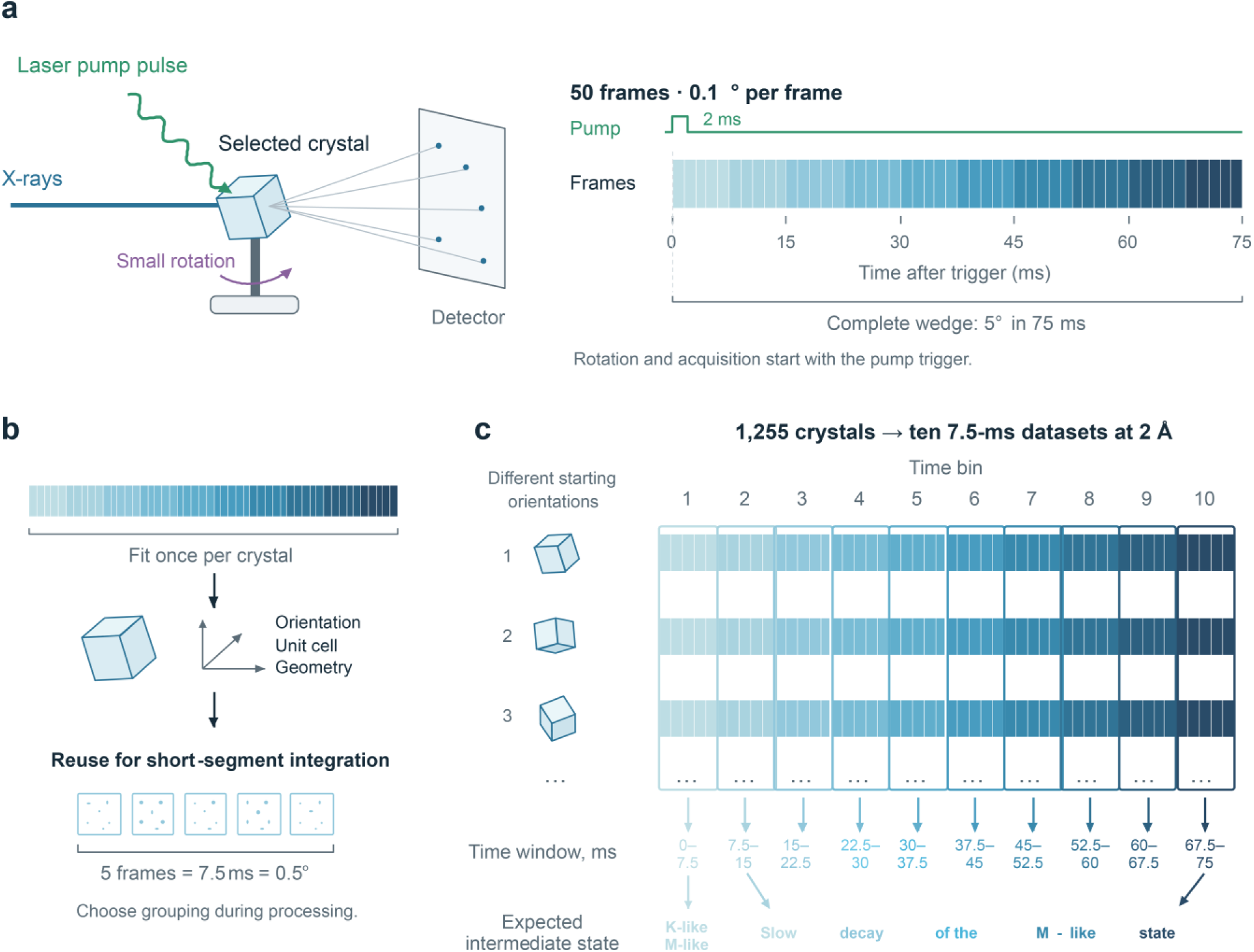
Serial small-wedge acquisition recovers a time series from one crystal ensemble. a, Diffraction images are recorded continuously during a small crystal rotation synchronized to a single activation pulse. In the illustrated acquisition, a 2-ms laser pulse begins at time zero, and 50 consecutive 1.5-ms frames span 0.1° each, producing a 5° wedge over 75 ms. b, Crystal orientation, unit-cell parameters and diffraction geometry are refined using the complete wedge and reused to integrate shorter image segments. Temporal binning is chosen during processing; in the example, five frames correspond to 7.5 ms and 0.5°. c, Segments covering the same time window are merged across crystals with different starting orientations. The same ensemble of 1,255 measured crystals yielded ten consecutive 7.5-ms datasets at 2 Å, spanning 0–75 ms. The state labels summarize the qualitative expectations from in-crystal spectroscopy (Supplementary Fig. 4): K-like and M-like populations in the first window, followed by slow decay of the M-like population.

The complete rotation wedge is first processed with XDS to determine the crystal orientation, unit-cell parameters and diffraction geometry. These parameters are then applied to the shorter temporal segments, which may contain too few reflections to determine their geometry precisely. Because reflections are recorded as they pass through the diffraction condition, conventional rotation-data integration can be used. Scaling used XSCALE. Correlation-based clustering with ccCluster identifies isomorphous wedges for segment merging.^17,23,24^

Time bins are defined retrospectively during processing. For consecutive frames with an exposure time t, a segment containing N frames spans Nt; when acquisition begins at the activation trigger, the nominal boundaries of segment j are (j − 1)Nt and jNt. Longer segments increase the number of observations and improve signal at the cost of temporal averaging, whereas shorter segments provide finer temporal sampling, provided that completeness and signal-to-noise ratio remain sufficient. The practical bin width therefore depends on diffraction quality, orientation coverage, crystal symmetry and the magnitude of the structural response.

This flexibility is particularly useful when the kinetics of the crystallized protein are incompletely characterized. An initial merge can be used to test for a reproducible light-induced signal, after which the same raw data can be repartitioned into finer or coarser time windows. If the resulting signal or resolution is insufficient, additional crystals can be measured in a subsequent session using the same acquisition protocol and merged with the existing data, provided that acquisition geometry is compatible and crystal isomorphism and excitation conditions are maintained. Extending the recorded time interval requires longer acquisitions, with the X-ray flux adjusted to keep the accumulated dose below the level at which radiation damage compromises the measurement. Here, the useful exposure range was assessed using segmented dark wedges, as described below.

### Implementation with standard multi-crystal rotation acquisition

We implemented this strategy on the ESRF MASSIF-3 beamline using the Mesh&Collect workflow to locate and centre crystals in sealed Mylar sandwiches (Fig. 2). BR_MT_ was selected because it forms well-diffracting, untwinned crystals and can be crystallized in lipidic cubic phase (LCP), which holds the crystals securely in place within the sandwich. Crystal dimensions varied from approximately 20 to 100 µm along each of the three axes. The X-ray beam had a full width at half maximum (FWHM) of approximately 15 × 15 µm, and the excitation spot was approximately 100 µm in diameter. Crystals remained embedded in their surrounding LCP throughout transfer and data collection, which helped to protect them from mechanical and osmotic damage.

**Figure 2.**
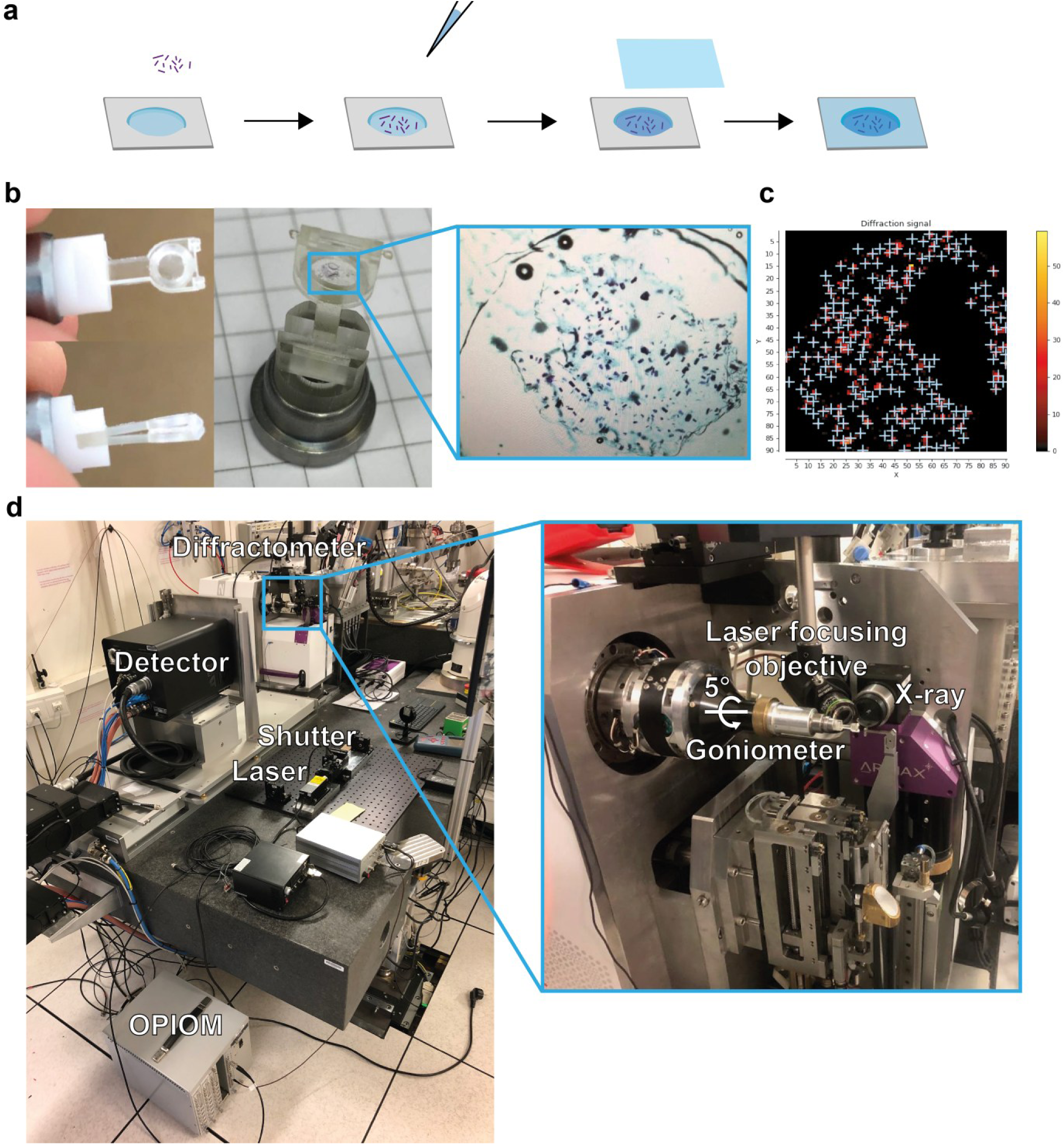
Mesh&Collect time-resolved implementation at MASSIF-3. a, LCP-grown crystals are sealed between Mylar films. b, The sandwich is mounted on an in-house 3D-printed holder with a magnetic base. c, A mesh scan locates diffracting crystals before wedge collection. d, A 532-nm laser, shutter and focusing objective are synchronized with the MD2 diffractometer and EIGER X 4M detector using OPIOM electronics. The approximately 100-µm excitation spot surrounds the X-ray beam (approximately 15 × 15 µm FWHM). The sample-delivery and synchronization implementation is distinct from the general acquisition principle in Fig. 1.

A diffractometer trigger synchronized the start of rotational data collection with a 2-ms pulse from a 532-nm laser. For each crystal, 50 images were collected at 0.1° per image, corresponding to a total rotation of 5°. With an exposure time of 1.5 ms per image, the acquisition spanned 75 ms. For these acquisition conditions, the estimated mean fluence-weighted absorbed dose increased from approximately 6.5 kGy in frames 1–10 to 28.5 kGy in frames 41–50, including prior mesh irradiation (Methods). Five consecutive images defined a 0.5° rotation segment acquired over 7.5 ms, yielding ten consecutive time windows from each accepted wedge (Table 1). Acquisitions at successive positions were separated by more than 3 s to allow recovery after possible excitation of neighbouring crystals.

**Table 1.** Acquisition accounting for the BR_MT_ benchmark.

| Parameter | Ground state | Early series | Extended series |
| --- | --- | --- | --- |
| Measured crystals | 349 | 1,255 | 790 |
| Wedges after clustering | 320 | 1,209 | 542 |
| Images per crystal | 50 | 50 | 50 |
| Exposure per image | 1.5 ms | 1.5 ms | 10 ms |
| Rotation per image / wedge | 0.1° / 5° | 0.1° / 5° | 0.1° / 5° |
| Pump duration | No pump | 2 ms | 2 ms |
| Nominal observation interval | Reference | 0–75 ms | 0–500 ms |
| Images per temporal bin | All 50 | 5 | 10 |
| Number of temporal bins | One reference | 10 | 5 |
| Time-bin width | Not applicable | 7.5 ms | 100 ms |
| Final wedges or segments per dataset | 320 | 1,200–1,205 | 424 first bin; 542 thereafter |
| Resolution | 1.8 Å | 2.0 Å | 2.0 Å |
| Mean DWD, frames 41–50 (kGy) | ~28.5 | ~28.5 | ~28.5 |

Counts refer to crystals measured as wedges, not all crystals loaded onto the supports. The three acquisitions total 2,394 measured crystals. Five additional dose-control datasets were generated by subdividing the ground-state wedges. Full data-collection and refinement statistics are in Supplementary Tables 1–4, 6 and 7. Mean diffraction-weighted dose (DWD) values are representative RADDOSE-3D estimates including prior mesh irradiation; they are averages over frames 41–50, not total or end-of-wedge doses. For the 10-ms series, transmission was reduced to 0.15 of that used for the 1.5-ms acquisitions, maintaining the same fluence per frame and modelled dose profile.

The setup combines a conventional diffractometer, a fast detector and established multi-crystal processing with additional laser excitation, shuttering, focusing optics and trigger electronics. It does not require optimization of a flowing sample stream: diffraction mapping locates usable crystals, which are then selected individually for rotation data collection. Mesh&Collect provides one practical implementation of the approach, which can in principle be transferred to other platforms capable of coordinating crystal positioning, small-angle rotation, excitation and detector timing.

### A membrane protein benchmark establishes structural sensitivity

We first collected 349 unilluminated wedges. Correlation-based clustering retained 320 wedges, yielding a 1.8 Å room-temperature ground-state dataset (PDB ID: 46NW) with 99.9% completeness and an outer-shell CC_1_/_2_ of 0.473 (Supplementary Table 1). The resulting maps resolved alternative conformations of the retinal chromophore, internal water molecules and alternative side-chain conformations around the proton-release region (Fig. 3). We also determined a cryogenic BR_MT_ ground-state structure at 1.48 Å resolution (PDB ID: 46NV), providing an independent reference for molecular replacement and structural comparison (Supplementary Table 2). Published room-temperature structures of wild-type bacteriorhodopsin (BR_WT_) provided additional context for interpreting the maps (Supplementary Fig. 3).

**Figure 3.**
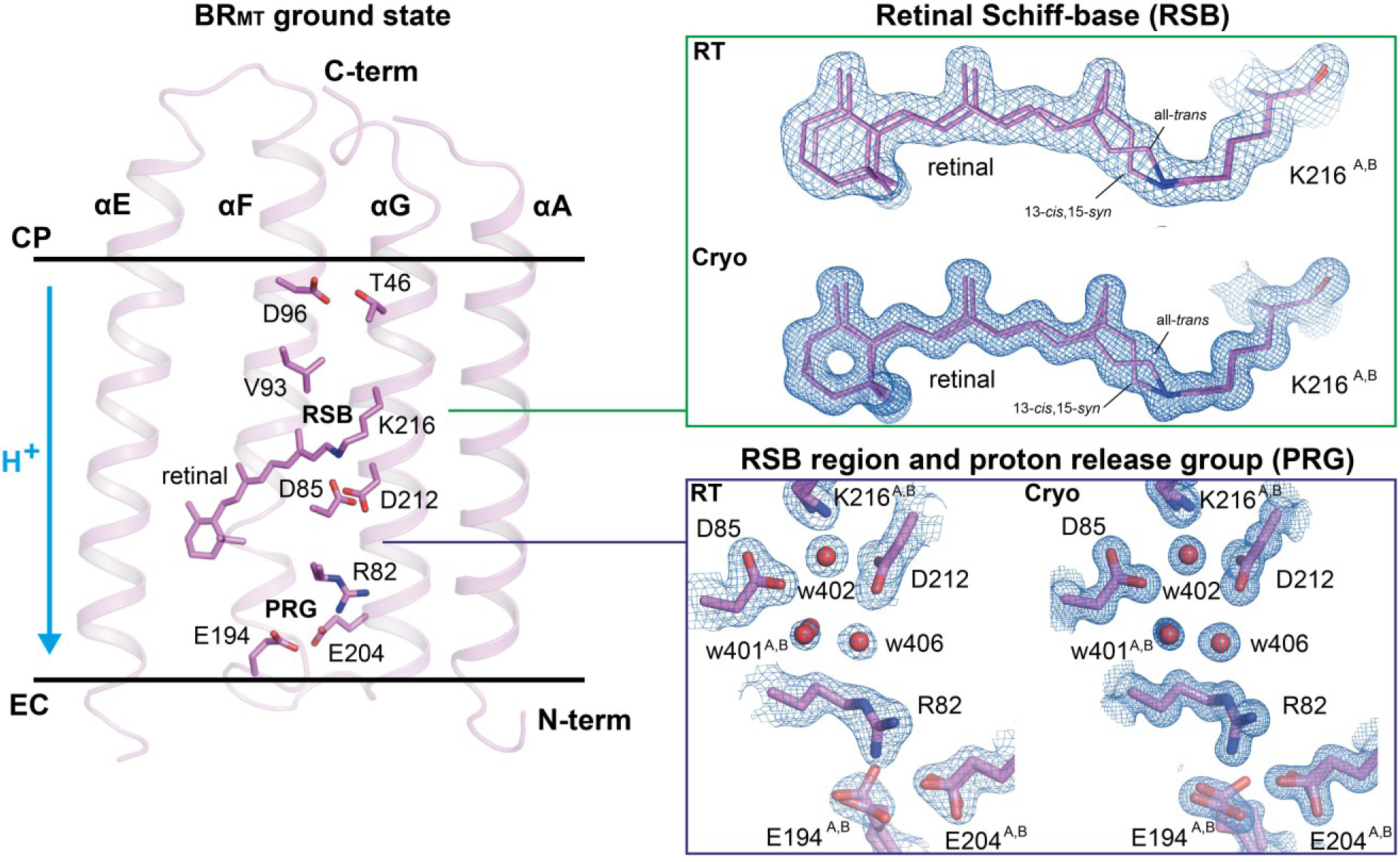
Room-temperature BR_MT_ ground-state structure at 1.8 Å resolution, with selected helices omitted for clarity. Key residues for proton pumping are depicted as sticks. The direction of proton pumping is indicated with a blue arrow. Insets compare the room-temperature model and the 1.48 Å cryogenic BR_MT_ reference determined in this study near the RSB and proton-release region. 2*m*Fo − *D*Fc maps are shown at 1σ. PPM-derived membrane boundaries are indicated; CP and EC denote the cytoplasmic and extracellular sides, respectively.^61^

Time-resolved *in crystallo* UV–Vis absorption spectroscopy was used to define the relevant observation window.^25^ Following nanosecond excitation at 532 nm, the early spectra contained red- and blue-shifted components consistent with K-like and M-like populations, respectively. The red-shifted component was detected from 10 µs to approximately 5 ms, after which the blue-shifted component dominated and decreased over the subsequent 500 ms (Supplementary Fig. 4a,b). Spectral traces at 415, 583 and 655 nm further resolved the evolution of these populations across the measured timescales (Supplementary Fig. 4c–e). These measurements motivated dense sampling at early times followed by coarser sampling over an extended observation window.

For the 75-ms diffraction series, we collected 1,255 wedges and retained 1,209 after clustering. Following segment integration and merging, between 1,200 and 1,205 segments contributed to each final time bin. All ten datasets extended to 2.0 Å resolution, with overall completeness of 98.9–99.9% and outer-shell CC_1_/_2_ values of 0.577–0.800 (Supplementary Table 3). Thus, the same ensemble of measured crystals supported the complete ten-window series. Including the separately collected ground-state dataset, a total of 1,604 crystals were measured for the ground-state reference and early time-resolved trajectory.

The 7.5–15-ms time window showed a clear light-induced structural signal (Fig. 4a). Difference maps and models refined against extrapolated structure-factor amplitudes revealed retinal isomerization from all-*trans* to 13-*cis,*15-*anti*, reorientation of R82 and reduced density for two water molecules near the retinal Schiff base (RSB). Together with the blue-shifted absorption, these changes are consistent with the M state reported for BR_WT_.^3,9,26^ Xtrapol8-generated maps guided modelling of an M-like conformation, which was combined with the ground-state conformation and refined against the observed structure-factor amplitudes (Supplementary Table 4).^27^ The room-temperature ground-state and M-like models retained a similar overall backbone conformation, with a Cα root-mean-square deviation (RMSD) of approximately 0.3 Å (Supplementary Fig. 5). Detection of difference density around retinal, side chains and internal water molecules in datasets with low activated-state occupancy demonstrates the structural sensitivity of the approach.

**Figure 4.**
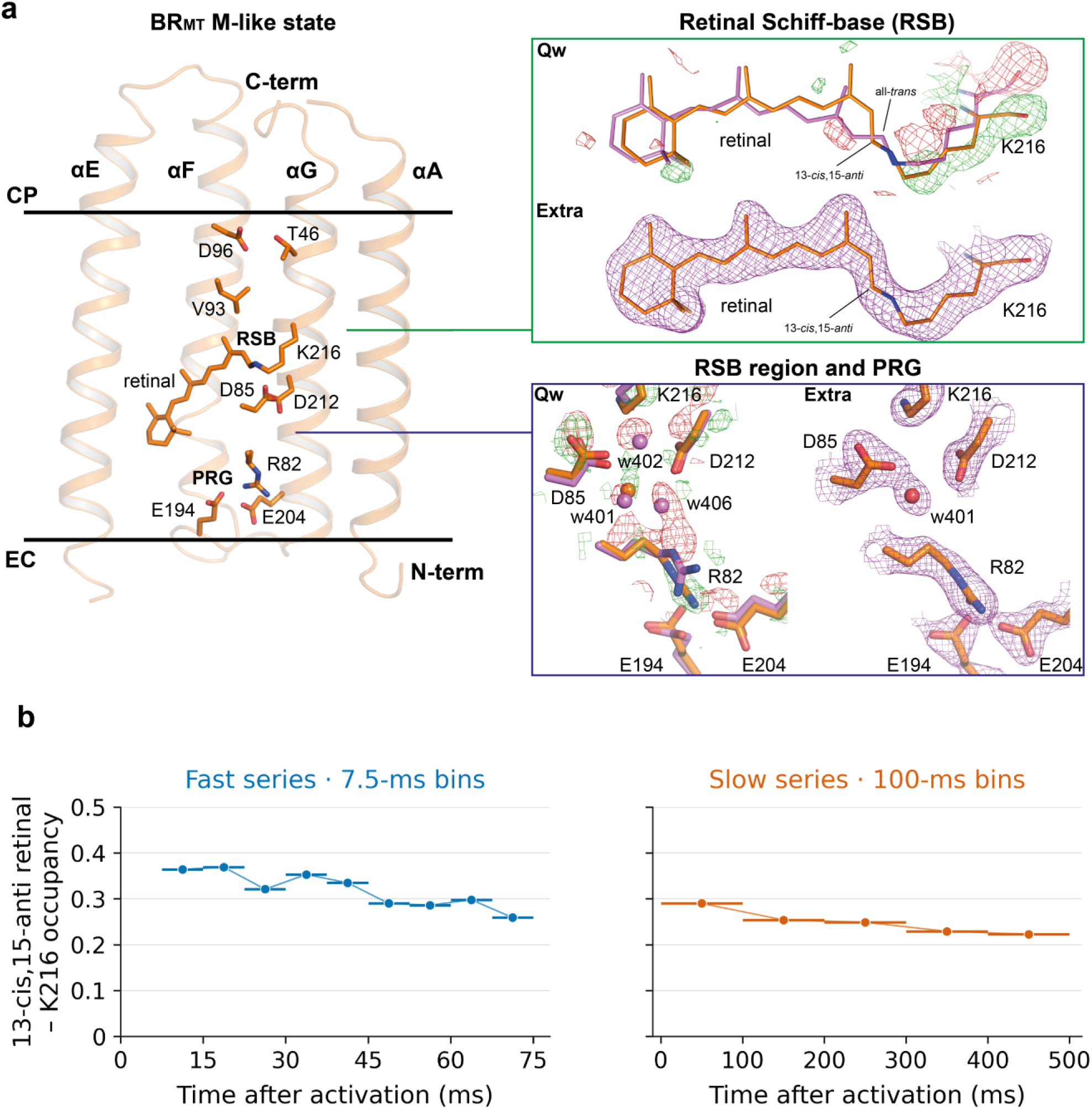
Repeated sampling resolves light-induced changes and their persistence. a, Room-temperature BR_MT_ M-like state structure, with selected helices omitted for clarity. Key residues for proton pumping are depicted as sticks. CP and EC denote the cytoplasmic and extracellular sides, respectively. Insets compare ground-state (violet) and M-like (orange) models near the RSB and proton-release region. Q-weighted observed-amplitude difference maps (Qw) and extrapolated maps (Extra) for the 7.5–15-ms dataset relative to the ground state are shown at 3σ and 1σ, respectively. b, Refined 13-*cis*,15-*anti* retinal–K216 occupancy (conformation D) for two independent acquisitions (Supplementary Table 5). Points mark bin midpoints and horizontal segments span the integration windows; connecting lines guide the eye. No kinetic fit or uncertainty estimate is implied. The fixed-population 0–7.5-ms model is omitted.

### Adjustable sampling captures early changes and slower relaxation

Occupancy refinement in phenix.refine against observed diffraction data, using the same ground-state and M-like conformations across successive time bins (Methods), revealed how the structural populations evolved. The refined occupancy of 13-*cis*,15-*anti* retinal decreased from 0.364 at 7.5–15 ms to 0.259 at 67.5–75 ms, with fluctuations between individual bins (Fig. 4b and Supplementary Fig. 6). Supplementary Table 5 lists the final model occupancies used for each plotted point. These values describe the structural model adopted for each time bin and should not be interpreted as independent measurements of kinetic rates.

The persistence of the structural signal motivated a second acquisition using 10-ms frames with the same 50-image, 5° wedge design. X-ray transmission was reduced in inverse proportion to the exposure time to maintain the same nominal total dose. Of 790 additional crystals, 542 passed clustering; the final merged datasets contained 424 segments in the first time window and 542 in each subsequent window. Grouping ten frames yielded five 100-ms bins spanning 0–500 ms, each reaching 2.0 Å resolution (Supplementary Table 6). The refined occupancy of 13-*cis*,15-*anti* retinal decreased from 0.290 in the 0– 100-ms bin to 0.223 in the 400–500-ms bin. Together, the two series demonstrate that acquisition duration and temporal bin width can be adapted to the observed structural response. The 0–75-ms and 0–500-ms series used separate crystal populations; within each series, the same ensemble contributed to successive time windows, avoiding the need for a new crystal population for each delay.

The spectral data suggest that the first 7.5-ms bin may contain a mixture of early and later photoproducts. A Q-weighted Fourier difference map comparing the 0–7.5-ms and 7.5–15-ms datasets revealed additional retinal features that were less apparent when the ground state alone was used as the reference (Supplementary Fig. 7). Guided by *in crystallo* spectroscopy and previous BR_WT_ structures, we tentatively assign the early component to a K-like state.^28^ The adjacent-bin comparison illustrates how dense temporal sampling can resolve evolving structural features even when individual bins contain mixed populations. Separate K-like (PDB ID: 46NX) and M-like (PDB ID: 46NY) models refined against extrapolated amplitudes are described in Supplementary Table 9. Both extrapolated models retain 13-*cis* retinal but differ in the torsional geometry of the retinal–Lys216 linkage and the conformation of Arg82. Differences in the modelled water sites around the RSB and the Glu194/Glu204 proton-release region further distinguish the K-like and M-like structures.

### Segmented dark wedges provide a dose-resolved control

Because each wedge records consecutive images of the same crystal, the dark data also provide an internal control for structural changes that accumulate with X-ray exposure. Radiation-induced changes in bacteriorhodopsin can occur at relatively low doses and potentially confound light-induced signals, making dose control important for synchrotron time-resolved measurements.^29,30^ We divided the unilluminated wedges into five ten-image segments and compared their merged electron-density maps. The estimated mean fluence-weighted absorbed doses for frames 1–10, 11–20, 21–30, 31–40 and 41–50 were approximately 6.5, 12.0, 17.5, 23.0 and 28.5 kGy, respectively, retaining all preceding irradiation, including the mesh scan (Methods). No prominent local difference features were observed near the RSB at the displayed contour levels (Supplementary Fig. 8 and Supplementary Table 7).

Diffraction statistics weakened across the dark series, indicating accumulating global radiation damage. Nevertheless, no apparent site-specific damage was detected near the RSB at the displayed contour levels, supporting the interpretation of the light-induced changes within the investigated dose range. The control cannot exclude damage incurred before or during the first segment or interactions between photoactivation and X-ray exposure, but it illustrates a practical advantage of the method: segmenting pilot dark wedges before time-resolved data collection allows the useful dose range to be assessed and acquisition settings optimized accordingly.

## Discussion

We obtained a complete ten-window millisecond trajectory of a membrane protein at 2.0 Å resolution from 1,255 microcrystals. Because a dose-limited exposure is distributed across fine-sliced diffraction images, each wedge samples all time windows, and time bins can be redefined retrospectively during processing. In total, 2,394 crystals were measured at room temperature: 1,255 for the early trajectory, 349 for the ground-state dataset and 790 for the extended observation window.

This approach complements existing time-resolved methods. XFEL experiments capture ultrafast structural snapshots using femtosecond X-ray pulses, but typically require many crystals to assemble a time series.^9,31^ At synchrotrons, still-image and burst-acquisition approaches can reach short delays without continuous crystal rotation during the probe, and repeated-frame measurements already capture multiple time delays from individual crystals.^3,5,6^ Small-wedge acquisition adds information on reflection profiles and diffraction geometry provided by rotation to this repeated temporal sampling.

In such experiments, sample use efficiency depends both on sample delivery and on the temporal information recovered from each illuminated crystal. Viscous extrusion is well suited to LCP-grown crystals, although stable flow requires control of the material’s rheological properties.^32^ Fixed targets and drop-on-demand delivery can reduce the loss of unused material, while pump–probe measurements must avoid unintended pre-illumination, particularly between neighbouring crystals on fixed targets.^33,34,35,36,37^ Reaction initiation must also match the delivery scheme; hit-and-return scheduling and liquid application provide established approaches for optical and ligand triggering, respectively.^38,39^

Recent developments further expand the options when sample is limited. Microsecond serial acquisition at the ESRF ID29 beamline resolved ligand binding to the A2A adenosine receptor from 2,000 merged frames in a static experiment,^40^ while a recent preprint reports ligand-triggered functional rotation crystallography from individual crystals over longer timescales.^41^ Serial small-wedge acquisition addresses a complementary regime, in which many small crystals each contribute a dose-limited sequence of diffraction images.

The number of crystals measured here is substantially smaller than the number of indexed single-shot images reported for several previous rhodopsin datasets. For example, a channelrhodopsin SFX study reported 16,442–24,015 indexed single-shot images per delay, compared to the 1,255 measured crystals supporting all ten time windows here.^15^ This difference illustrates that fewer crystals may be required for a time-resolved trajectory, but does not constitute a direct comparison of total sample consumption. The experiments differ in target, crystal size, resolution, dose regime and activation fraction. Indexed images do not account for material lost during sample delivery, and other approaches can also reuse crystals across multiple delays (Supplementary Table 8). What the present approach demonstrates is that a single ensemble of 1,255 crystals supports the complete early trajectory.

For membrane proteins and other difficult-to-crystallize targets, obtaining a structural trajectory from a limited crystal ensemble may considerably expand the range of feasible experiments. Individual selection of usable crystals avoids the requirement for a dense suspension optimized for delivery, while established rotation-data processing and the ability to add compatible wedges from subsequent sessions make the experiment adaptable to the available sample. Existing small-wedge datasets for GPCRs and time-resolved GPCR photopharmacology provide natural starting points for extending the approach beyond BR_MT_.^8,16,17^

The approach has several limitations. Temporal resolution depends on the activation-pulse duration, the X-ray exposure time and the time intervals over which reflection intensities are integrated. During rotation, each reflection diffracts over a finite angular range, its rocking curve, which, at a given rotation speed, determines when and for how long it contributes to the recorded images. Different reflections can therefore weight different portions of the same time bin, so the nominal 7.5-ms bin width alone does not fully describe the temporal response.^17,20^ Dose accumulation limits the useful trajectory length, and weak diffraction may require broader bins or additional crystals.^29,30^ Absolute dose estimates are also limited by variation in incident flux and crystal dimensions. Heterogeneous activation and incomplete recovery must also be considered, particularly for slowly reversible or irreversible reactions, and excitation of neighbouring crystals may need to be controlled.^37,42^ Because the spectroscopic measurements used nanosecond excitation whereas the diffraction experiments used a 2-ms pump pulse, the resulting populations are not expected to be identical. When multiple intermediates coexist, structural occupancies and assignments of early states remain model-dependent. Models derived from extrapolated amplitudes additionally depend on the selected extrapolation fraction and on uncertainties in the underlying difference amplitudes. These effects can be evaluated using complementary spectroscopy, dose controls and dedicated excitation tests.

The present implementation targets millisecond-to-second dynamics, a timescale relevant to many functional transitions in membrane proteins.^3,4,5,6,7^ Here, temporal resolution was limited mainly by the available laser power for activation and by the EIGER detector readout rate. Faster excitation, detectors and X-ray gating could extend the temporal range of the approach.^40^ In particular, nanosecond excitation and faster detectors, such as JUNGFRAU,^43^ could enable temporal sampling approaching 1 ms, provided that sufficient diffraction signal is retained. The approach’s immediate value, however, lies in making efficient use of limited crystal material: a single crystal ensemble supports a complete time series, temporal binning can be adapted during processing, acquisition settings can be tuned to the tolerable dose, and the method is compatible with established multi-crystal rotation workflows.

## Methods

### Protein expression and purification

BR_MT_ was expressed in *Escherichia coli* using a modified BR_WT_ expression protocol.^44^ The native halobacterial nucleotide sequence contained two silent substitutions, C9A and G12A, corresponding to residues A2 and Q3. A C-terminal GSGIEGRSGAPHHHHHHHH tag included a factor Xa cleavage site and an affinity tag. The construct was cloned into pSCodon 2.1 (Delphi Genetics) through NdeI and XhoI sites; T17A, T24A and T47A were introduced by PCR-based site-directed mutagenesis.

E. coli BL21(DE3) cells carrying the construct were grown at 37 °C in 2-l baffled flasks containing ZYP-5052 autoinduction medium and 100 µg ml^−1^ ampicillin, with shaking at 120 r.p.m. At OD_600_ = 1.0, the temperature was reduced to 20 °C, and 10 µM all-*trans* retinal and additional ampicillin were added. Ampicillin was added again at OD_600_ = 3.0. After 14 h of incubation, cells were harvested and disrupted in an M-110P homogenizer (Microfluidics). Membranes were isolated by ultracentrifugation at 90,000 × g for 1 h at 4 °C, resuspended in 20 mM Tris-HCl, pH 8.0, and 100 mM NaCl, and solubilized overnight with 1% (w/v) n-dodecyl-β-D-maltoside (DDM).

Insoluble material was removed by centrifugation at 90,000 × g for 1 h at 4 °C. Imidazole was added to the supernatant to 10 mM before loading onto Ni-NTA resin (Qiagen). The column was washed with five column volumes of 50 mM sodium phosphate, pH 8.0, 100 mM NaCl, 50 mM imidazole and 0.2% DDM, followed by one column volume of the same buffer at pH 6.0. Protein was eluted in 50 mM sodium phosphate, pH 6.0, 100 mM NaCl, 0.5 M imidazole and 0.3% DDM, concentrated with a 30-kDa-cutoff Amicon Ultra device (Millipore), and loaded onto a 30-ml Superdex 200 Increase column (GE Healthcare) equilibrated in 50 mM sodium phosphate, pH 6.0, 100 mM NaCl and 0.2% DDM. The coloured protein peak was pooled. The yield was approximately 1–3 mg of purified protein per litre of culture.

### Crystallization and sample mounting

For *in meso* crystallization, we followed workflows similar to those used in our previous studies of OLPVR1 and MAR.^45,46,47^ Protein concentrated to 40 mg ml^−1^ was mixed with monoolein (Nu-Chek Prep), pre-melted at 42 °C, at a 3:2 monoolein-to-protein-solution ratio using 100-µl Hamilton syringes. An NT-8 robot (Formulatrix) dispensed 150 nl of LCP and 400 nl of precipitant into each well of 96-well glass plates (Paul Marienfeld). The precipitant contained 2.5 M ammonium sulfate and 0.1 M sodium acetate, pH 5.2. Approximately 50–150 cubic crystals per well appeared within one month with dimensions ranging from approximately 20 to 100 µm along each axis.

Crystals were transferred together with surrounding transparent LCP onto 10-µm Mylar film for crystallography or 100-µm cyclic olefin copolymer (COC) film for spectroscopy. Pre-cut wells in double-sided adhesive tape (467MP, 3M, for crystallography; LCP-100, SWISSCI, for spectroscopy) defined the sample chamber. Before sealing with a second film, 1 µl of a solution containing 15% (v/v) glycerol, 2.5 M ammonium sulfate and 0.1 M sodium acetate, pH 5.2, was added. Glycerol reduces dehydration of LCP during room-temperature handling.^48^

### Time-resolved absorption spectroscopy

Spectra of BR_MT_ crystals were recorded at the ESRF time-resolved in crystallo optical spectroscopy (TR-icOS) setup.^25^ The COC sandwich was aligned in three dimensions to obtain sufficient absorbance near 560 nm while avoiding detector saturation and minimizing background. A xenon flash-lamp module (L11316-11, Hamamatsu) and AvaSpec-ULS2048CL-EVO-RS-UA spectrophotometer (Avantes) recorded absorption before excitation and at defined delays from 10 µs to 500 ms after a nanosecond pulse from a tunable Surelite EX laser (Amplitude Technologies). Spectra were levelled by subtracting the mean absorbance between 800 and 900 nm using the icOS toolbox.^49^ Difference spectra were calculated by subtracting the pre-excitation spectrum.

Excitation was at 532 nm with a nominal fluence of 50 mJ cm^−2^. Pre-excitation spectra were recorded before successive pulses to monitor recovery and persistent spectral changes. Excitation conditions influence rhodopsin photochemistry,^50^ and spectra collected with nanosecond pulses were used to identify relevant timescales rather than to impose identical populations on the millisecond-pumped diffraction data.

### Rotation acquisition and synchronization

The data-collection and processing concept used here was originally described in ref.^51^ Diffraction was recorded at room temperature at the ESRF ID30A-3 (MASSIF-3) beamline.^22^ Data were collected in two sessions for the room-temperature ground state (30 June 2023 and 21 May 2024), three for the 0–75-ms series (19 July 2023, 24 April 2024 and 26 April 2024), and two for the 0–500-ms series (18 January 2024 and 9 February 2024). A Mylar film sandwich was attached to an in-house 3D-printed magnetic-base sample holder on an MD2 diffractometer (Arinax), with the film plane initially perpendicular to the X-ray beam (approximately 15 × 15 µm FWHM). The Mesh&Collect workflow used Dozor software scores to localize and rank diffracting positions.^21^ Positions showing diffraction beyond 3 Å were selected for 5° wedges, recorded as 50 images at 0.1° per image on an EIGER X 4M detector (Dectris).

A 532-nm laser (CNI) was coupled through a fast mirror-based shutter (Standa) and a 600-µm optical fibre to a 20× objective (Olympus). An OPIOM digital input/output module (ESRF) received the diffractometer trigger also used to initiate detector acquisition and generated an inverted high signal of 2 ms to open the laser shutter. Time zero denotes this acquisition and activation trigger. With 1.5-ms frames, the nominal observation interval was 0–75 ms; for the extended series, 10-ms frames covered 0–500 ms. Increasing the exposure from 1.5 to 10 ms was compensated by reducing the X-ray transmission to 1.5/10 = 0.15 of that used for the 1.5-ms acquisition. This preserved the incident fluence per frame and the nominal total dose for each 50-image wedge.

For laser alignment, a pinhole was centred at the beam and rotation centre. A 532-nm LED (Thorlabs) allowed coarse alignment of the focusing objective. A photodiode (Thorlabs) downstream of the pinhole then provided a signal for a two-dimensional alignment scan followed by position optimization along three motorized axes. The excitation spot was approximately 100 µm in diameter, estimated with the on-axis viewer, and the measured power was 4 mW (Nova II, Ophir). A 2-ms pulse therefore delivered approximately 8 µJ; for a uniform circular 100-µm spot this corresponds to a nominal incident fluence of approximately 100 mJ cm^−2^. This geometrical estimate does not include the beam profile, reflection, scattering or absorption by the films and LCP; consequently, the local fluence at individual crystals may differ from the nominal incident value.^42^

Ground-state mesh scans and wedges were acquired without laser excitation. For time-resolved acquisitions, crystal positions were localized with the laser off; the laser was enabled before collection of the triggered wedges. The interval between acquisitions at successive positions exceeded 3 s. Table 1 summarizes the acquisition series and their separately measured crystal populations.

### Absorbed-dose estimation

Absorbed doses were estimated using RADDOSE-3D (v. 5.0.1067).^52^ Crystals were modelled as centred 50 × 50 × 50 µm cuboids irradiated by a Gaussian beam with a full width at half maximum of 15 × 15 µm at 12.812 keV. The unit-cell composition was derived from the refined protein–retinal structure, with an estimated solvent fraction of 0.527. The solvent contained 15% (v/v) glycerol, 2.5 M ammonium sulfate and 0.1 M sodium acetate, pH 5.2; its density was assumed to be 1.20 g ml^−1^.

The modelled acquisition comprised 50 consecutive 1.5-ms exposures over a total rotation of 5°, at an incident flux of 3.82 × 10^11^ photons s^−1^. This value represented the mean flux used for the calculations; fluxes between measurements varied from approximately 3 × 10^11^ to 6 × 10^11^ photons s^−1^. Prior mesh irradiation was represented by a local 5 × 5 grid centred on the collection position, with a 15-µm step, 1.5-ms exposure per position and threefold higher flux, thereby accounting for overlapping beam tails. Calculations used 1-µm voxels and 0.01° angular increments during rotation, without modelling photoelectron escape, fluorescence escape or diffraction-intensity decay.

Mean fluence-weighted absorbed doses (the “Average Diffraction Weighted Dose” output with DDM Simple) were calculated separately for each ten-frame subset while retaining all preceding irradiation. The estimated doses for frames 1–10, 11–20, 21–30, 31–40 and 41–50 were approximately 6.5, 12.0, 17.5, 23.0 and 28.5 kGy, respectively. With the transmission scaled inversely with exposure time and the same prior mesh irradiation, the same modelled dose sequence applies to corresponding ten-frame subsets of the 0–500-ms series. Crystal dimensions varied from approximately 20 to 100 µm along each of the three axes. Together with the variation in incident flux, this limits the precision of the calculation: the reported values are representative model-based estimates rather than exact doses for individual crystals.

### Integration, clustering and temporal segmentation

An in-house wrapper applied XDS and XSCALE to the small wedges, following established multi-crystal processing strategies.^16,23^ An initial run used a 3 Å cutoff without a prescribed space group or unit cell to establish orientation and geometry. Wedges were reintegrated with refined parameters and the predominant crystal symmetry, to 1.8 Å for the ground state or 2.0 Å for time-resolved data. Pairwise intensity correlations of full wedges were analysed with ccCluster to select isomorphous wedges (Supplementary Fig. 9).^24^ Initial clustering was performed on complete wedges before temporal segmentation and thus selected for overall data isomorphism rather than activation homogeneity within individual time bins. The processing workflow calls its clustering-distance threshold “CC vicinity”; a value of 0.6 was used for whole-wedge selection. An additional segment-level clustering step was required for the 0–100-ms dataset, as described below.

For the ground-state merge, 320 of 349 wedges passed the selected threshold. Thresholds of 0.4, 0.6 and 0.8 were compared with the complete merge; 0.6 gave the highest outer-shell CC_1_/_2_ among the tested settings and was therefore selected for subsequent model building (Supplementary Table 1). For the 75-ms series, 1,209 of 1,255 wedges passed clustering. Each was split into ten five-image segments, reintegrated with the geometry obtained from its parent wedge and merged with corresponding segments from other crystals. Segments that could not be integrated successfully were excluded. Consequently, 1,200–1,205 segments contributed to each final time-bin merge (Supplementary Table 3). The common 2 Å cutoff retained an outer-shell CC_1_/_2_ greater than 0.5 for every bin.

For the 500-ms series, 542 of 790 wedges passed initial whole-wedge clustering. Ten-image segments generated five 100-ms windows. The initial 0–100-ms merge of 542 segments had an overall CC_1_/_2_ of 0.104 (10.4%) and was therefore reclustered with ccCluster at a CC vicinity of 0.5. This retained 424 segments and yielded an overall CC_1_/_2_ of 0.992 (0.552 in the highest-resolution shell); 542 segments were retained for each subsequent window (Supplementary Table 6).

To assess radiation damage, the dark dose series was produced by dividing the ground-state wedges into five ten-image segments and reintegrating to 2 Å. These merges contained 308–309 successfully integrated segments (Supplementary Table 7).

### Cryogenic reference structure

The cryogenic BR_MT_ ground-state reference was determined at 1.48 Å using data collected at ESRF ID23-1 at 100 K. Crystals were grown using the same monoolein LCP crystallization conditions as for the time-resolved measurements, including 2.5 M ammonium sulfate and 0.1 M sodium acetate, pH 5.2. Before opening each well, 100 µl of precipitant supplemented with 15% (v/v) glycerol was added for cryoprotection. Crystals were harvested in MiTeGen micromounts, flash-cooled and stored in liquid nitrogen. Diffraction images were processed with XDS, while STARANISO was used for anisotropic truncation and scaling.^23,53,54^ The anisotropic diffraction limits were 1.48, 1.68 and 1.65 Å along a*, b* and c*, respectively. Molecular replacement with MOLREP used a BR_WT_ ground-state model (PDB ID: 7Z0A).^55^ The model was rebuilt in Coot, refined with phenix.refine using four translation–libration–screw (TLS) groups, with occupancies fixed, and validated with MolProbity.^56,57,58^ This separately measured reference provided an independent structural comparison with the room-temperature model; its statistics are reported in Supplementary Table 2.

### Model building and structural comparison

The cryogenic BR_MT_ ground-state structure served as the reference model to build the room-temperature structure (Supplementary Table 2). For molecular replacement of the room-temperature dataset with Phaser, the model was reduced to a single polyalanine conformation.^59^ The solution was rebuilt with AutoBuild, iteratively adjusted in Coot and refined with phenix.refine; geometry was assessed using MolProbity.^56,57,58,60^ Room-temperature ground-state and time-resolved refinement statistics are given in Supplementary Tables 4, 6 and 7. Membrane boundaries shown in structural illustrations were predicted with the PPM server.^61^ Cα RMSDs were calculated by a single least-squares superposition of all corresponding Cα atoms, without outlier rejection.

Xtrapol8 (v. 1.2.9) was used to calculate Q-weighted difference and extrapolated maps for the 7.5–15-ms dataset using the room-temperature ground-state dataset as reference.^27^ Calculations used 12–2 Å data and isotropic B-factor scaling. Candidate extrapolation fractions f were scanned from 0.10 to 0.85 in steps of 0.05, with an extrapolation factor α = 1/f. The selected f = 0.30 (α = 3.33) maximized the sum of selected integrated difference-density peaks; the Pearson map correlation was 0.827, close to its maximum of 0.828 at f = 0.25. Maps extrapolated at f = 0.30 guided an M-like conformation, which was combined with a ground-like conformation. The final two-state models contained retinal–K216 in alternate conformations A and D; a separate 13-*cis*,15-*syn* ground-state retinal component was not included. Refinement against observed diffraction data was done in phenix.refine. For the 7.5–75-ms and 0–500-ms series, individual coordinates, rigid-body placement and occupancies of alternate conformations A and D were refined over five macrocycles, without individual B-factor refinement. Coordinate refinement accommodated the merged unit-cell differences. Avoiding individual B-factor refinement reduces occupancy–displacement correlation, but the resulting occupancies remain conditional on this structural model.

For the 0–7.5-ms dataset, maps calculated against the ground state were dominated by features resembling the M-like model. We therefore also calculated Q-weighted difference and extrapolated maps using the 7.5–15-ms dataset as reference, applying the comparative difference-map strategy used previously for mixed ligand states in cytochrome P450.^62^ Features preferentially present in the first bin guided an additional K-like conformation. For this comparison, f was scanned from 0.05 to 0.40 in steps of 0.05. The adopted f = 0.20 (α = 5.0) coincided with the maximum Pearson map correlation (CC = 0.794), whereas difference-map maximization selected f = 0.10 (α = 10.0). These criterion-dependent estimates describe the inter-bin contrast rather than an independently measured absolute K-like population. In the final three-conformation model, ground-state (A), K-like (C) and M-like (D) occupancies were held at 0.50, 0.20 and 0.30, respectively, during coordinate and individual isotropic B-factor refinement.

For the dose control, observed-amplitude difference maps compared each later dark segment with the first dark segment. Maps were calculated in Phenix and inspected at the contour levels shown in Supplementary Fig. 8.

## Data availability

The structural datasets comprise the room-temperature ground state, ten 7.5-ms time bins, five 100-ms time bins, five dark dose-control bins, the 1.48 Å cryogenic BR_MT_ reference, and separately refined K-like and M-like extrapolated-data models. PDB accession codes are 46NV (cryogenic ground state), 46NW (room-temperature ground state), 46NX (extrapolated K-like model) and 46NY (extrapolated M-like model); 46NZ, 46OA, 46OB, 46OC, 46OD, 46OE, 46OF, 46OG, 46OH, 46OI (ten 7.5-ms bins, in chronological order); 46OJ, 46OK, 46OL, 46OM, 46ON (five 100-ms bins, in chronological order); and 46OO, 46OP, 46OQ, 46OR, 46OS (five dark dose-control bins, in increasing dose order). Raw and processed diffraction data and source spectroscopy data will be made publicly available upon publication. Dataset statistics are provided in Supplementary Tables 1–4, 6, 7 and 9. The occupancies plotted in Fig. 4b and Supplementary Fig. 6 are listed in Supplementary Table 5.

## Code availability

Diffraction data were processed with XDS, XSCALE and ccCluster, coordinated by an in-house wrapper. The in-house wrapper is available upon request.

## Acknowledgements

We thank the staff of the ESRF and EMBL Hamburg, especially Daniele de Sanctis, Shibom Basu, Julien Orlans, David Flot, Kirill Kovalev and Gleb Bourenkov, for helpful discussions, and Gordon Leonard for supporting the project. We appreciate the help of Peter van der Linden (ESRF) with printing sample holder on the 3D printer. S.B. acknowledges support from the ESRF PhD programme (2020–2022). This work was supported by ANR-19-CE11-0026 and ANR-25-CE11-1470.

## Author contributions

S.B., V.G. and I.M. conceived the project. D.B., C.B. and T.B. expressed the protein. R.A. crystallized the protein. N.C., S.E. and A.R. collected and processed time-resolved spectroscopy data from crystals. I.M. supervised instrumentation. S.B. supervised diffraction data collection, processing and structure refinement. A.P. assisted with instrumentation, data collection and processing. S.B. and V.G. wrote the manuscript with contributions from all authors. C.M.-D., M.S.-L., and E.B. supported the project and helped with the manuscript refinement.

## Competing interests

The authors declare no competing interests.

## Notes

### Competing Interest Statement

The authors have declared no competing interest.

