## Supplementary Information for "Sample-efficient time-resolved X-ray crystallography with serial small wedges"

Sergei Bukhdruker and coauthors

### Structural interpretation of the bacteriorhodopsin benchmark

The T17A/T24A/T47A triple mutant of bacteriorhodopsin (BR_MT_) was used to test whether small-wedge temporal sampling could detect local structural changes in a membrane protein. Wild-type bacteriorhodopsin is denoted BR_WT_ throughout. Photocycle assignments draw on in-crystal absorption measurements and established structures. Spectroscopic kinetics can differ between crystals and membranes,^1^ and structural labels do not imply that the crystalline mutant reproduces every step of the BR_WT_ cycle.

The room-temperature ground-state model contains all-*trans* and 13-*cis*,15-*syn* retinal populations at approximately 70% and 30%, respectively. The coexistence of these two isomers is a working hypothesis, and their identities and relative occupancies remain tentative at the present resolution. Modelling is further complicated by the absence of inter-dataset difference electron-density maps analogous to those used to guide the K-like and M-like models. Definitive characterization would require dedicated structural validation, ideally with diffraction data approaching 1 Å resolution, as illustrated by high-resolution studies of OLPVR1 and archaerhodopsin-3.^2,3^ Such validation is beyond the scope of this method-focused study.

In the 7.5–15-ms light-activated dataset, the additional conformation includes 13-*cis*,15-*anti* retinal, reorientation of R82 toward E194/E204 and reduced occupancy of waters w402 and w406. These observations are consistent with an M-like population. They do not independently determine proton positions or establish individual proton-transfer events at 2 Å resolution. The Cα root-mean-square deviation is approximately 0.28 Å between ground-state and M-like models and approximately 0.26 Å between room-temperature and 1.48 Å cryogenic ground-state models.

Under its own excitation conditions, spectroscopy detected a red-shifted component on timescales overlapping the first diffraction window. Comparison of the first diffraction window with the next supports a K-like model with a distorted retinal Schiff base (RSB) linkage, a smaller R82 reorientation and shifted waters near the RSB (Supplementary Fig. 7). The retinal distortion is consistent with the K state reported for BR_WT_.^4^ The descriptor 15-twisted denotes a C15=Nζ geometry displaced from ideal 15-*syn* or 15-*anti* configurations, consistent with the terminology of prior K-state analysis.^5^ The nominal three-state populations (0.50/0.20/0.30) were held fixed during the final coordinate and B-factor refinement; they are not refined occupancies. Different population changes can contribute to the inter-bin difference, so the comparison does not uniquely deconvolve a K state.

The persistent R82 population in the extended series and the absence of clearly resolved later spectral components are observations, not proof of a direct M-to-ground-state pathway. Low occupancy, spectral overlap, the 100-ms bins and crystal-lattice effects can conceal other intermediates. The data likewise do not identify R82 reorientation as a rate-limiting step. These mechanistic questions require finer kinetic sampling and independent validation beyond the methodological demonstration presented here.

### Interpreting crystal and diffraction counts across studies

Supplementary Table 8 summarizes selected published rhodopsin datasets. The reported counts describe the observations or crystal contributions used in individual merges, as labelled in the underlying studies. They should not be read as total sample consumption. Multiple patterns can originate from one crystal, time windows can share a crystal population and unused material can be lost during delivery. The asterisk identifies source entries explicitly noted as collecting several time windows from the same crystal; its absence does not prove a one-pattern-per-crystal relationship.

The room-temperature acquisitions comprise 349 measured crystals for the ground state, 1,255 for the 75-ms series and 790 for the 500-ms series: 2,394 crystals in total. The 1.48 Å cryogenic reference was collected separately (Supplementary Table 2). In the channelrhodopsin comparison, 16,442–24,015 indexed single-shot images per delay correspond to 13.1–19.1 times the 1,255 measured crystals supporting our full ten-window series (Supplementary Table 8). This compares reported diffraction contributions rather than total sample volume. A matched benchmark also requires protein mass, crystal dimensions, hit and indexing rates, activation fraction, dose, completeness, resolution and the number of reconstructed windows.

### Supplementary Figure 1

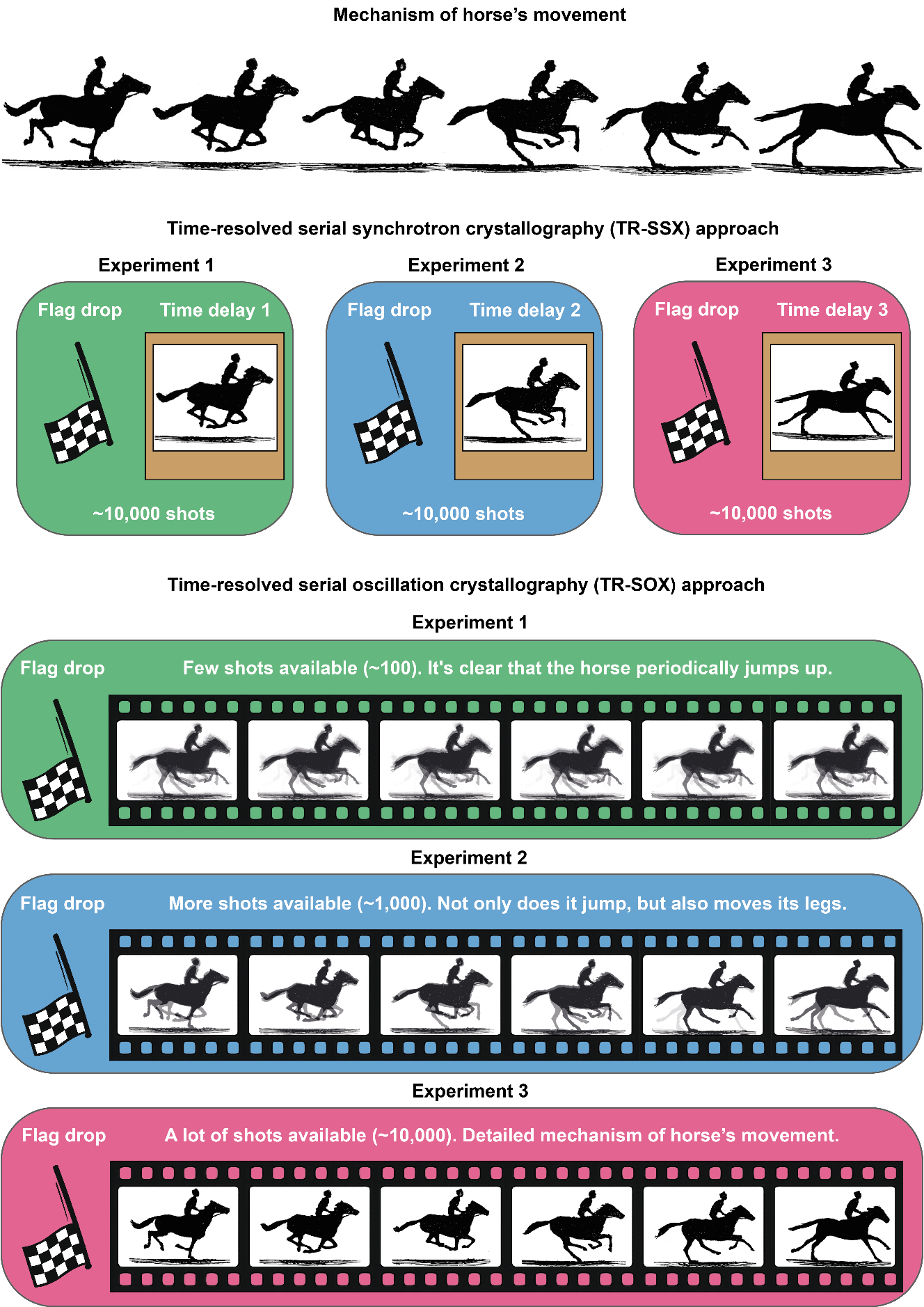

Supplementary Figure 1. Visual analogy for repeated temporal sampling. Successive horse positions represent reaction progress. The upper example allocates independent snapshots to selected delays; the lower example illustrates how recording a sequence from each specimen can contribute to multiple windows, with sampling adjusted during analysis. Shot counts are illustrative and are not measured crystal requirements or universal benchmarks. Modern still-image burst methods can also record several delays per crystal. The small-wedge strategy combines repeated temporal sampling with rotation information; its quantitative performance is reported in the main text.

### Supplementary Figure 2

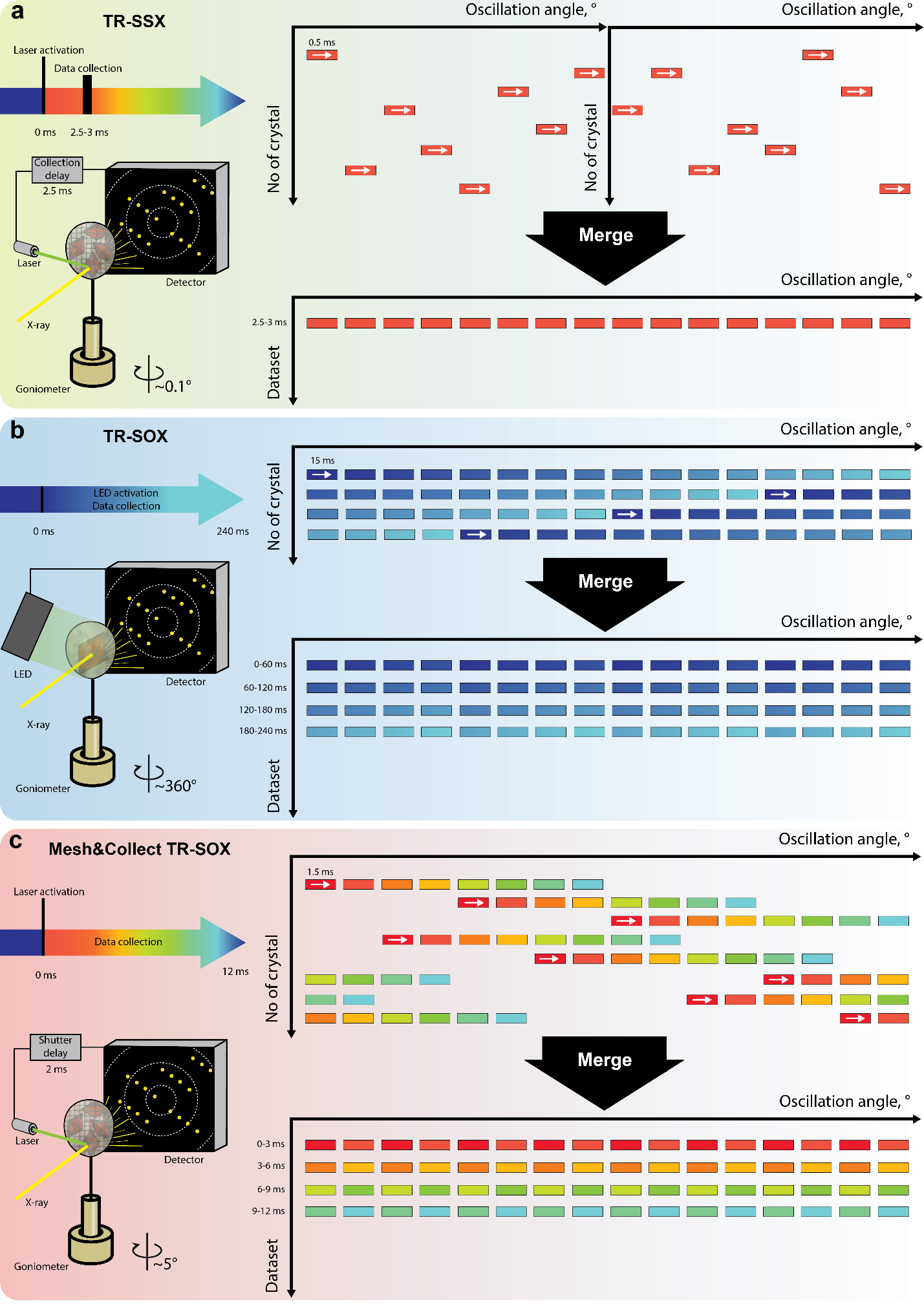

Supplementary Figure 2. Comparison of time-resolved acquisition schemes. a, Single-delay serial still acquisition. b, Continuous-illumination time-resolved serial oscillation acquisition. c, Pulse-triggered short-wedge acquisition illustrated with Mesh&Collect. Timelines show how image sequences are grouped into temporal windows. The numerical timings and angular ranges are schematic examples; the measured protocols in this study are specified in Fig. 1 and Table 1. The 0.1° symbol in a indicates narrow orientation sampling, not intentional rotation during a still exposure. Bin widths describe sampling intervals and do not include all contributions to the temporal response.

### Supplementary Figure 3

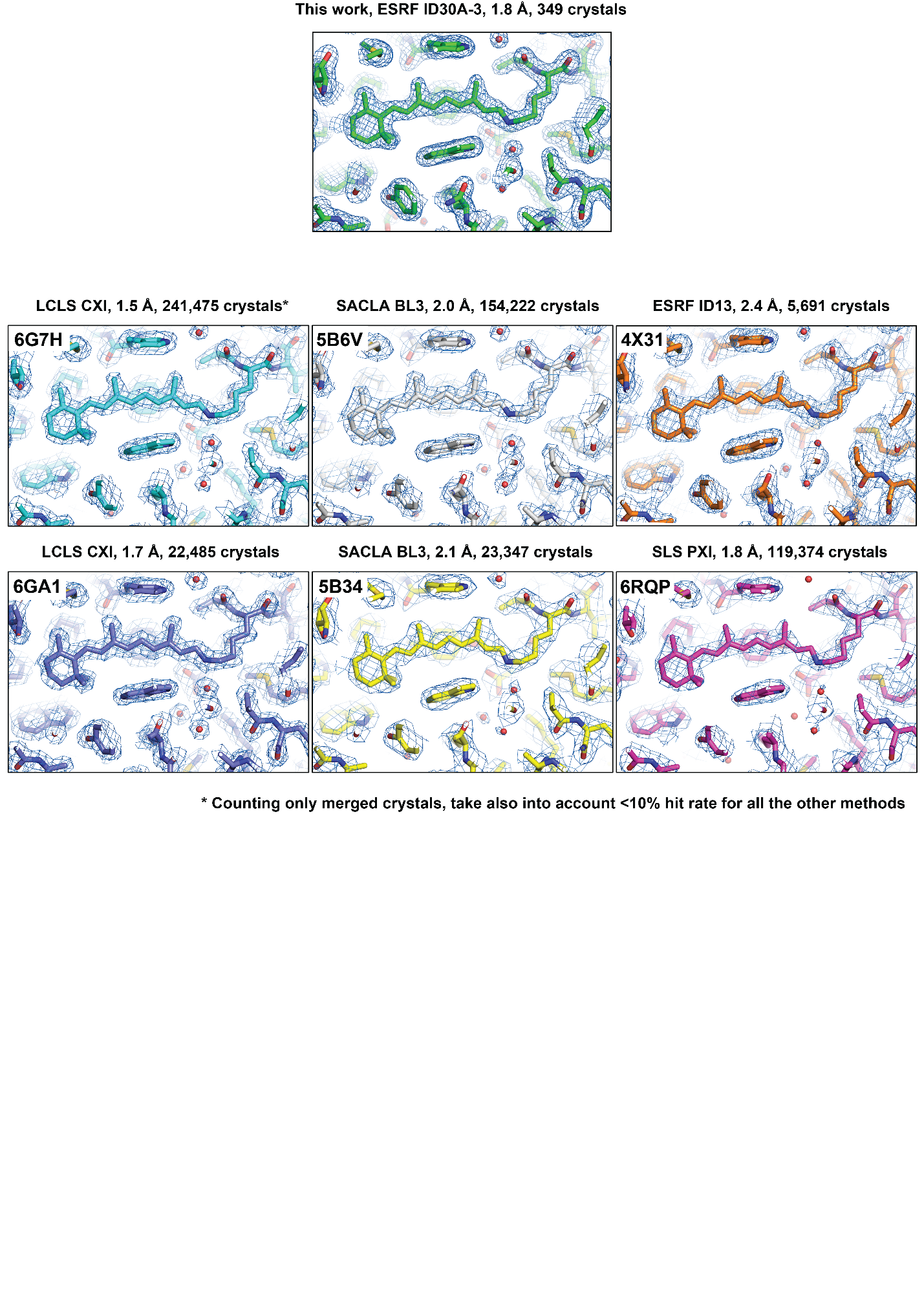

Supplementary Figure 3. Room-temperature ground-state map comparison. Published BR_WT_ ground-state models and the BR_MT_ model are compared using 2mFo − DFc maps contoured at 1.3σ. Only conformation A is shown for BR_MT_. These panels illustrate map features across different studies; resolution, crystal form, sample history and data processing were not matched, so they do not establish relative acquisition efficiency. Counts labelled as crystals in the published-study panels describe merge inputs and are not total sample consumption.

### Supplementary Figure 4

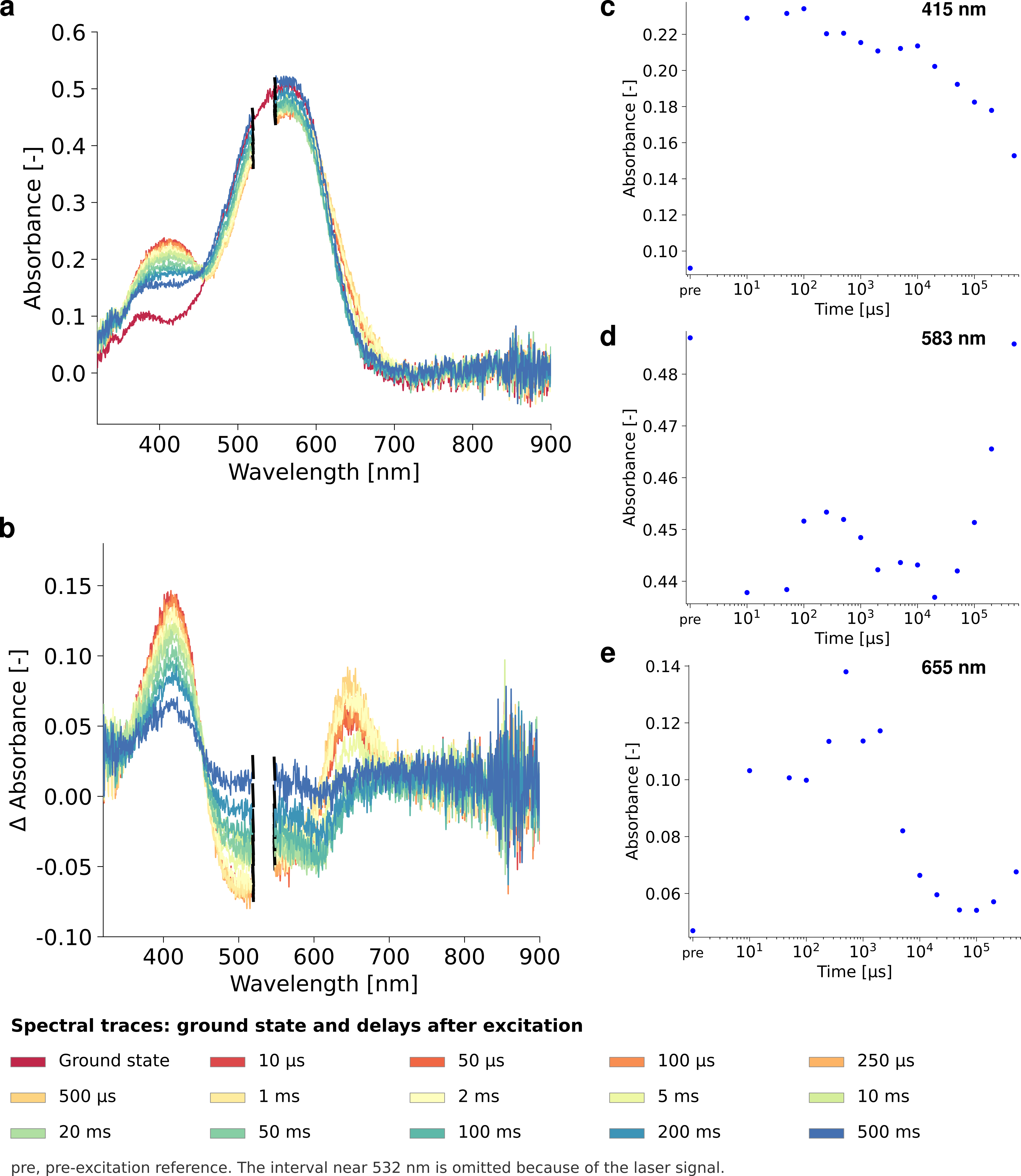

Supplementary Figure 4. Time-resolved absorption spectroscopy of BR_MT_ crystals. a, Pre-excitation ground-state spectrum (dark red) and spectra recorded at the delays indicated in the legend after nanosecond 532-nm excitation. b, Corresponding post-excitation-minus-pre-excitation difference spectra. The interval around 532 nm is omitted because of the excitation-laser signal. c–e, Absolute absorbance at 415, 583 and 655 nm, respectively. The label “pre” identifies the pre-excitation reference at the left; positive post-excitation delays are shown on a logarithmic scale. The 415-nm signal remains above the reference at 500 ms, so recovery of the ground-state absorbance at this wavelength is incomplete within the plotted interval.

### Supplementary Figure 5

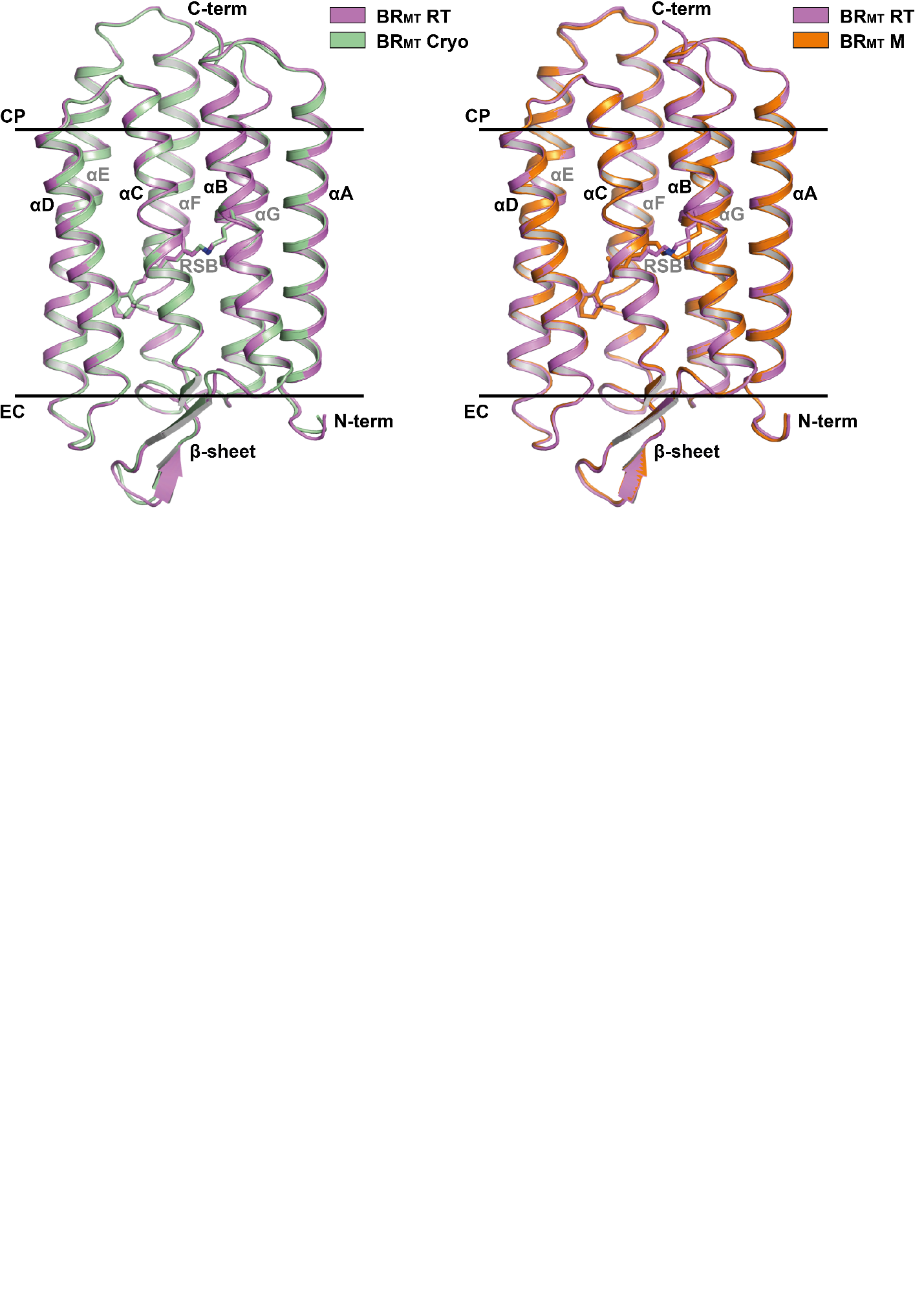

Supplementary Figure 5. Alignment of BR_MT_ models. The 1.48 Å cryogenic ground-state reference determined in this study, room-temperature ground state and M-like BR_MT_ models are shown in green, violet and orange. Retinal is shown as sticks; PPM-predicted membrane boundaries are black.^6^ Reported Cα root-mean-square deviations are approximately 0.26 Å between the two ground-state models and 0.28 Å between room-temperature ground and M-like models. These comparisons include all corresponding Cα atoms. The N terminus and β-sheet region show additional local differences in the displayed alignment.

### Supplementary Figure 6

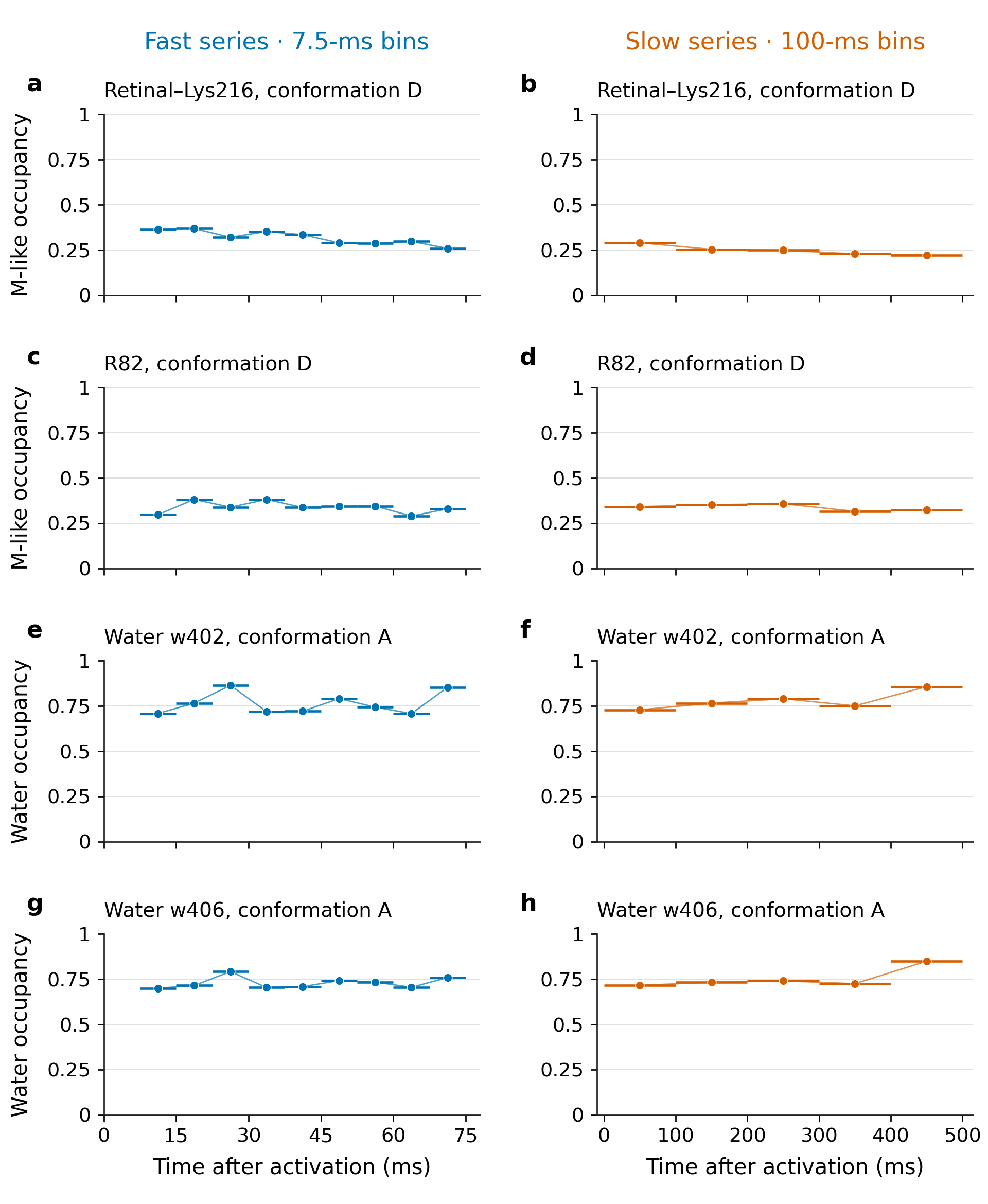

Supplementary Figure 6. Refined structural-marker occupancies from the final models. Rows show retinal–K216 (conformation D), R82 (conformation D), w402 (HOH A507, conformation A) and w406 (HOH A506, conformation A). Columns show the independent 75-ms and 500-ms acquisitions. Points lie at bin midpoints; horizontal segments denote integration windows and connecting lines guide the eye. No kinetic fit or uncertainty estimate is implied. The fixed-population 0–7.5-ms model is excluded. Every plotted value is listed in Supplementary Table 5.

### Supplementary Figure 7

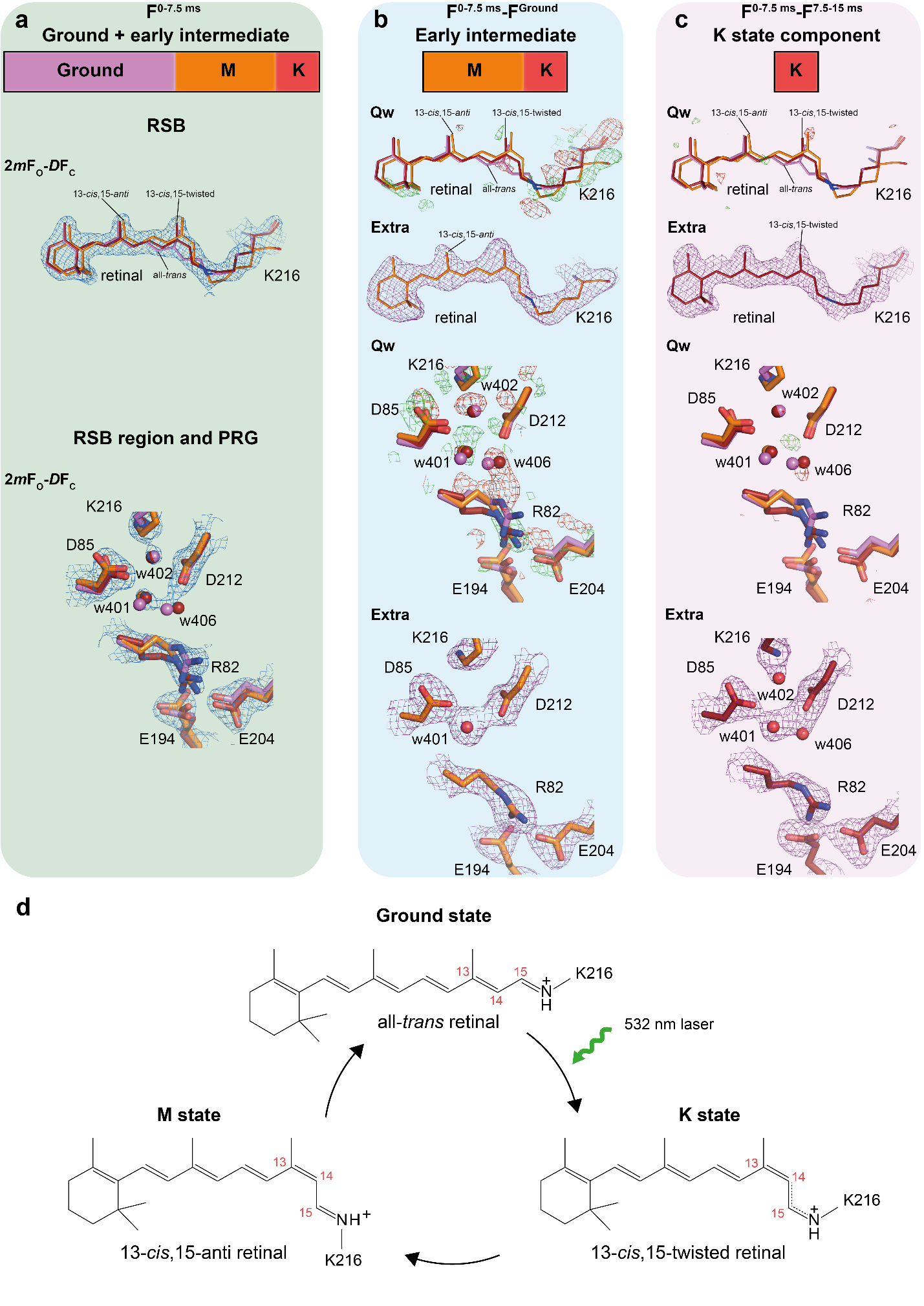

Supplementary Figure 7. Temporal contrast in the early mixed population. a, The 0–7.5-ms model contains ground-state, K-like and M-like populations fixed at 50%, 20% and 30% during the final refinement. 2*m*Fo − *D*Fc maps are shown at 1σ. b, Difference and extrapolated maps against the ground state emphasize features resembling the M-like component. c, Comparison with the 7.5–15-ms dataset emphasizes changes between the two mixed populations. Q-weighted difference maps are shown at 3σ and extrapolated maps at 1σ. The labels “K state component” and “K state” denote a provisional K-like assignment. “15-twisted” describes a distorted C15=Nζ geometry.^5^

### Supplementary Figure 8

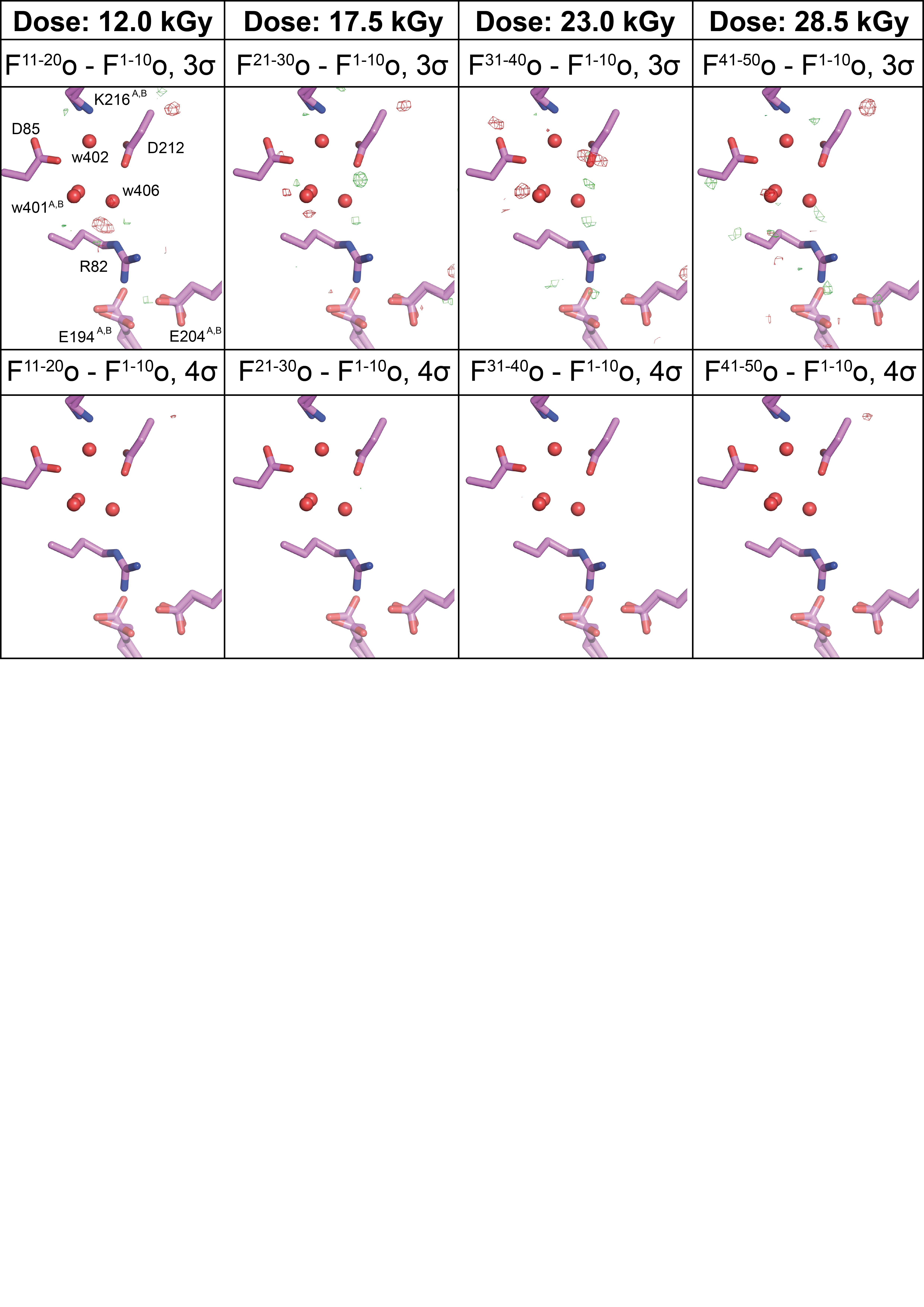

Supplementary Figure 8. Dose-dependent differences in unilluminated BR_MT_. Fifty-frame dark wedges were divided into five ten-frame segments with estimated mean diffraction-weighted doses of approximately 6.5, 12.0, 17.5, 23.0 and 28.5 kGy, respectively, retaining all preceding irradiation, including the mesh scan. These are subset-averaged doses, not dose increments or end-of-subset doses. The maps compare each later segment with the first (images 1–10). Residues near the RSB are shown as sticks and waters as spheres. Observed-amplitude difference maps are contoured at 3σ and 4σ. The absence of prominent local features at these thresholds does not exclude smaller changes or damage before or during the first segment.

### Supplementary Figure 9

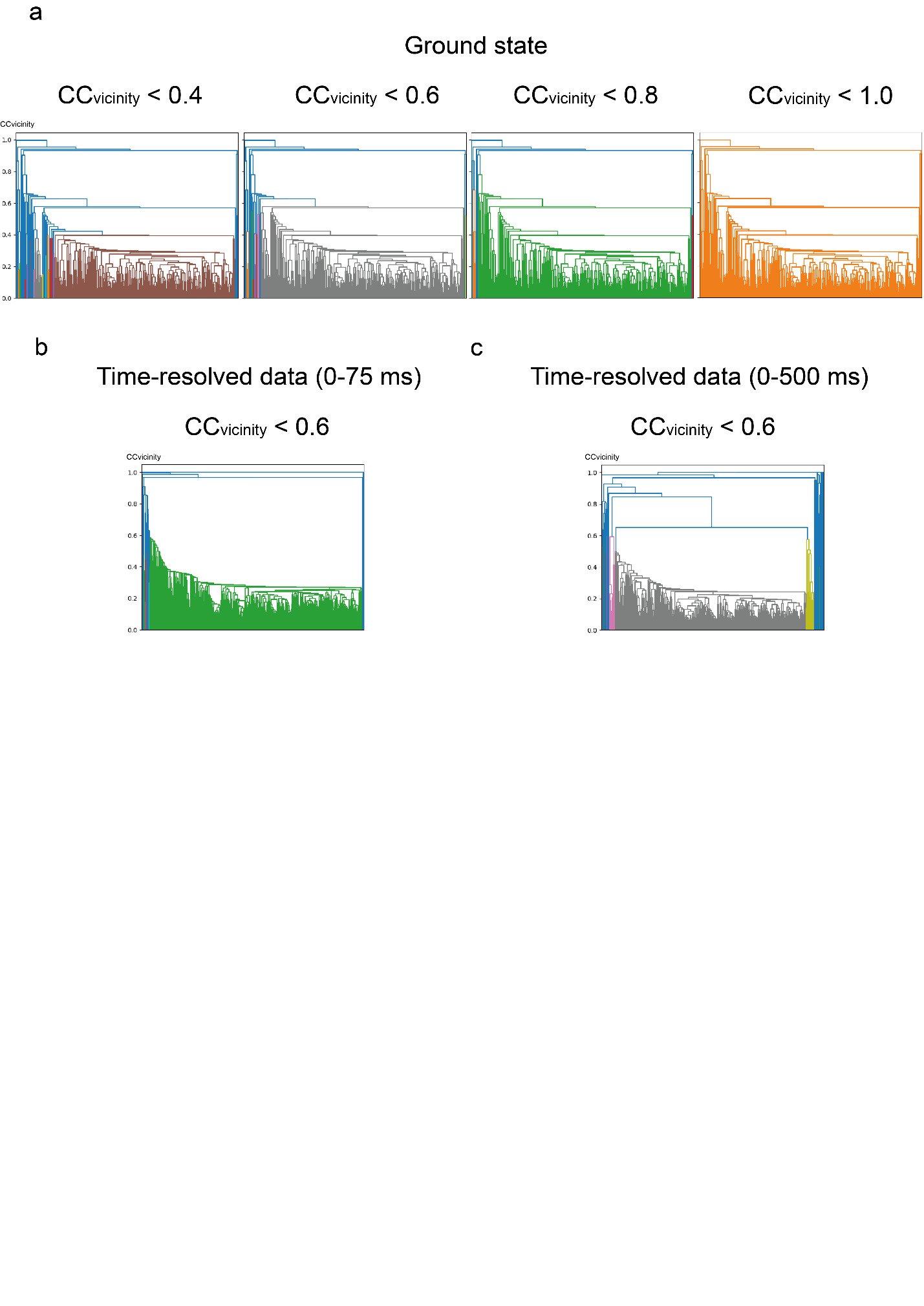

Supplementary Figure 9. Clustering of crystal wedges before temporal segmentation. Intensity correlations between wedges were analysed using ccCluster.^7^ Panels show the ground-state acquisition, the 0–75-ms acquisition and the 0–500-ms acquisition. A clustering-distance threshold (“CC vicinity”) of 0.6 was selected using the ground-state comparison and applied to the time-resolved series. The 0–100-ms segments underwent an additional clustering step at CC vicinity 0.5, retaining 424 of the 542 segments selected by the initial whole-wedge clustering (Methods).

### Supplementary Table 1

Ground-state data collection and the effect of the clustering threshold.

| **Isomorphism cutoff criteria for merging** | **All small-wedge datasets merged** | **CC vicinity < 0.8** | **CC vicinity < 0.6 (Best)** | **CC vicinity < 0.4** |
| --- | --- | --- | --- | --- |
| Small-wedge datasets merged/collected | 349/349 | 336/349 | 320/349 | 284/349 |
| Total number of images used | 17,450 | 16,800 | 16,000 | 14,200 |
| Space group | C222 | C222 | C222 | C222 |
| Unit cell |  |  |  |  |
| a, Å | 117.47 | 117.47 | 117.47 | 117.47 |
| b, Å | 120.29 | 120.29 | 120.29 | 120.29 |
| c, Å | 36.81 | 36.81 | 36.81 | 36.81 |
| Resolution, Å | 30.00 – 1.80 (1.85 – 1.80)* | 30.00 – 1.80 (1.85 – 1.80) | 30.00 – 1.80 (1.85 – 1.80) | 30.00 – 1.80 (1.85 – 1.80) |
| Total number of observations/unique | 1,556,109/24,740 (100,229/1,806) | 1,508,025/24,740 (97,971/1,806) | 1,440,661/24,742 (94,231/1,807) | 1,278,609/24,742 (83,967/1,807) |
| Completeness, % | 99.9 (99.9) | 99.9 (99.9) | 99.9 (99.9) | 99.9 (99.9) |
| Multiplicity | 63 (55) | 61 (54) | 58 (52) | 52 (46) |
| <(I)/σ(I)> | 11.1 (0.3) | 11.0 (0.3) | 11.0 (0.4) | 10.9 (0.4) |
| CC(1/2) | 0.998 (0.451) | 0.998 (0.468) | 0.998 (0.473) | 0.996 (0.459) |
| Rmeas | 0.463 (5.712) | 0.403 (5.671) | 0.366 (5.634) | 0.300 (5.403) |

Values in parentheses refer to the highest-resolution shell. All datasets are in space group C222 with α = β = γ = 90°.

### Supplementary Table 2

Data collection and refinement of the cryogenic BR_MT_ ground-state reference.

| **Dataset** | **Cryogenic ground state** |
| --- | --- |
| Data collection statistics | |
| Images taken | 1–900 |
| Total number of images used | 900 |
| Space group | C222 |
| Unit cell |  |
| a, Å | 116.01 |
| b, Å | 119.58 |
| c, Å | 36.10 |
| Resolution, Å | 37.70 – 1.48 (1.64 – 1.48) |
| Anisotropic resolution limits, Å | 1.48 / 1.68 / 1.65 (a*, b*, c*) |
| Total number of observations/unique | 203,847/30,486 (10,020/1,524) |
| Spherical completeness, % | 71.7 (14.1) |
| Ellipsoidal completeness, % | 89.8 (48.9) |
| Multiplicity | 6.7 (6.6) |
| <(I)/σ(I)> | 5.9 (1.5) |
| CC(1/2) | 0.995 (0.461) |
| Rmeas | 0.227 (2.387) |
| Refinement statistics | |
| PDB ID | 46NV |
| Resolution, Å | 37.70 – 1.48 |
| Rwork/Rfree | 0.1833/0.2142 |
| No. reflections work/free | 28,986/1,491 |
| No. of atoms |  |
| Protein | 1,889 |
| Retinal | 40 |
| Solvent | 87 |
| Lipids | 216 |
| B-factors, Å² |  |
| Protein | 15.94 |
| Retinal | 9.62 |
| Solvent | 27.13 |
| Lipids | 37.98 |
| R.m.s. |  |
| Bonds, Å | 0.006 |
| Angles, ° | 0.882 |
| Ramachandran favoured | 100.00 |
| Ramachandran outliers | 0 |

Values in parentheses refer to the highest-resolution shell. Completeness is spherical after anisotropic truncation; ellipsoidal completeness is 89.8% (48.9%). Space group C222; α = β = γ = 90°.

### Supplementary Table 3

Data collection for the ten consecutive 7.5-ms bins.

| **Segment** | **1** | **2** | **3** | **4** | **5** |
| --- | --- | --- | --- | --- | --- |
| Images taken | 1–5 | 6–10 | 11–15 | 16–20 | 21–25 |
| Time window, ms | 0–7.5 | 7.5–15 | 15–22.5 | 22.5–30 | 30–37.5 |
| Segments merged/collected | 1,203/1,255 | 1,202/1,255 | 1,202/1,255 | 1,203/1,255 | 1,205/1,255 |
| Total number of images used | 6,015 | 6,010 | 6,010 | 6,015 | 6,025 |
| Space group | C222 | C222 | C222 | C222 | C222 |
| Unit cell |  |  |  |  |  |
| a, Å | 117.21 | 117.22 | 118.32 | 118.92 | 118.15 |
| b, Å | 117.79 | 117.95 | 120.27 | 121.53 | 119.96 |
| c, Å | 36.47 | 36.49 | 36.81 | 36.99 | 36.77 |
| Resolution, Å | 30.00 – 2.00 (2.05 – 2.00)* | 30.00 – 2.00 (2.05 – 2.00) | 30.00 – 2.00 (2.05 – 2.00) | 30.00 – 2.00 (2.05 – 2.00) | 30.00 – 2.00 (2.05 – 2.00) |
| Total number of observations/unique | 307,413/17,559 (23,076/1,279) | 310,945/17,618 (23,955/1,297) | 320,160/18,230 (23,330/1,321) | 322,893/18,460 (18,902/1,199) | 320,577/18,182 (24,291/1,342) |
| Completeness, % | 99.8 (99.5) | 99.9 (100) | 99.7 (99.4) | 98.9 (86.6) | 99.9 (99.9) |
| Multiplicity | 18 (18) | 18 (18) | 18 (18) | 17 (16) | 18 (18) |
| <(I)/σ(I)> | 6.9 (1.3) | 9.8 (1.9) | 9.5 (1.6) | 9.5 (1.3) | 9.5 (1.6) |
| CC(1/2) | 0.993 (0.694) | 0.997 (0.800) | 0.996 (0.712) | 0.994 (0.584) | 0.995 (0.742) |
| Rmeas | 0.308 (1.522) | 0.232 (1.415) | 0.237 (1.577) | 0.243 (1.594) | 0.241 (1.561) |

Values in parentheses refer to the highest-resolution shell. All datasets are in space group C222 with α = β = γ = 90°.

### Supplementary Table 3 continued

Data collection for the ten consecutive 7.5-ms bins.

| **Segment** | **6** | **7** | **8** | **9** | **10** |
| --- | --- | --- | --- | --- | --- |
| Images taken | 26–30 | 31–35 | 36–40 | 41–45 | 46–50 |
| Time window, ms | 37.5–45 | 45–52.5 | 52.5–60 | 60–67.5 | 67.5–75 |
| Segments merged/collected | 1,204/1,255 | 1,203/1,255 | 1,204/1,255 | 1,200/1,255 | 1,201/1,255 |
| Total number of images used | 6,020 | 6,015 | 6,020 | 6,000 | 6,005 |
| Space group | C222 | C222 | C222 | C222 | C222 |
| Unit cell |  |  |  |  |  |
| a, Å | 117.64 | 118.65 | 118.35 | 117.17 | 118.54 |
| b, Å | 118.72 | 121.06 | 120.37 | 117.81 | 120.73 |
| c, Å | 36.59 | 36.93 | 36.84 | 36.47 | 36.90 |
| Resolution, Å | 30.00 – 2.00 (2.05 – 2.00) | 30.00 – 2.00 (2.05 – 2.00) | 30.00 – 2.00 (2.05 – 2.00) | 30.00 – 2.00 (2.05 – 2.00) | 30.00 – 2.00 (2.05 – 2.00) |
| Total number of observations/unique | 315,729/17,828 (23,454/1,294) | 321,188/18,438 (20,649/1,305) | 321,500/18,287 (23,429/1,323) | 310,008/17,572 (23,434/1,281) | 321,537/18,394 (21,974/1,338) |
| Completeness, % | 99.8 (99.9) | 99.5 (94.8) | 99.9 (99.9) | 99.9 (100) | 99.7 (98.4) |
| Multiplicity | 18 (18) | 17 (16) | 18 (18) | 18 (18) | 17 (16) |
| <(I)/σ(I)> | 9.7 (1.7) | 9.4 (1.2) | 9.4 (1.3) | 9.5 (1.6) | 9.2 (1.3) |
| CC(1/2) | 0.997 (0.742) | 0.997 (0.599) | 0.996 (0.691) | 0.996 (0.749) | 0.995 (0.577) |
| Rmeas | 0.233 (1.585) | 0.243 (1.703) | 0.254 (1.730) | 0.246 (1.585) | 0.256 (1.773) |

Values in parentheses refer to the highest-resolution shell. All datasets are in space group C222 with α = β = γ = 90°.

### Supplementary Table 4

Refinement statistics for the ground state and the 0–75-ms series.

| **Time window, ms** | **Dark** | **0–7.5** | **7.5–15** | **15–22.5** | **22.5–30** | **30–37.5** |
| --- | --- | --- | --- | --- | --- | --- |
| Intermediate state | Ground | K-like-M-like | M-like | Slow decay of M-like | | |
| PDB ID | 46NW | 46NZ | 46OA | 46OB | 46OC | 46OD |
| Resolution, Å | 29.37 – 1.80 | 29.30 – 2.00 | 29.31 – 2.00 | 29.58 – 2.00 | 29.73 – 2.01 | 29.54 – 2.00 |
| Rwork/Rfree | 0.1725/0.2006 | 0.1812/0.2278 | 0.1860/0.2284 | 0.1797/0.2082 | 0.1860/0.2188 | 0.1800/0.2090 |
| No. reflections work/free | 23,462/1,191 | 16,709/836 | 16,767/839 | 17,340/874 | 17,449/882 | 17,295/870 |
| No. of atoms |  |  |  |  |  |  |
| Protein | 1,873 | 5,160 | 3,438 | 3,438 | 3,438 | 3,438 |
| Retinal | 40 | 60 | 40 | 40 | 40 | 40 |
| Solvent | 50 | 99 | 67 | 67 | 67 | 67 |
| Lipids | 216 | 216 | 216 | 216 | 216 | 216 |
| B–factors, Å² |  |  |  |  |  |  |
| Protein | 35.87 | 34.94 | 32.92 | 32.92 | 32.92 | 32.92 |
| Retinal | 25.40 | 25.01 | 22.98 | 22.98 | 22.98 | 22.98 |
| Solvent | 47.24 | 40.32 | 43.48 | 43.48 | 43.48 | 43.48 |
| Lipids | 65.63 | 69.59 | 62.39 | 62.39 | 62.39 | 62.39 |
| R.m.s. |  |  |  |  |  |  |
| Bonds, Å | 0.015 | 0.002 | 0.002 | 0.003 | 0.003 | 0.004 |
| Angles, ° | 1.381 | 0.501 | 0.457 | 0.592 | 0.552 | 0.616 |
| Ramachandran favoured | 98.65 | 99.10 | 99.55 | 99.55 | 99.10 | 99.10 |
| Ramachandran outliers | 0 | 0 | 0 | 0 | 0 | 0 |

All datasets are in space group C222 with α = β = γ = 90°.

### Supplementary Table 4 continued

Refinement statistics for the ground state and the 0–75-ms series.

| **Time window, ms** | **37.5–45** | **45–52.5** | **52.5–60** | **60–67.5** | **67.5–75** |
| --- | --- | --- | --- | --- | --- |
| Intermediate | Slow decay of M-like | | | | |
| PDB ID | 46OE | 46OF | 46OG | 46OH | 46OI |
| Resolution, Å | 29.41 – 2.00 | 29.66 – 2.00 | 29.59 – 2.00 | 29.29 – 2.00 | 29.64 – 2.00 |
| Rwork/Rfree | 0.1806/0.2180 | 0.1863/0.2166 | 0.1832/0.2101 | 0.1946/0.2271 | 0.1814/0.2184 |
| No. reflections work/free | 16,963/847 | 17,448/887 | 17,389/881 | 16,723/836 | 17,437/887 |
| No. of atoms |  |  |  |  |  |
| Protein | 3,438 | 3,438 | 3,438 | 3,438 | 3,438 |
| Retinal | 40 | 40 | 40 | 40 | 40 |
| Solvent | 67 | 67 | 67 | 67 | 67 |
| Lipids | 216 | 216 | 216 | 216 | 216 |
| B–factors, Å² |  |  |  |  |  |
| Protein | 32.92 | 32.92 | 32.92 | 32.92 | 32.92 |
| Retinal | 22.98 | 22.98 | 22.98 | 22.98 | 22.98 |
| Solvent | 43.48 | 43.48 | 43.48 | 43.48 | 43.48 |
| Lipids | 62.39 | 62.39 | 62.39 | 62.39 | 62.39 |
| R.m.s. |  |  |  |  |  |
| Bonds, Å | 0.002 | 0.002 | 0.003 | 0.002 | 0.004 |
| Angles, ° | 0.500 | 0.524 | 0.589 | 0.431 | 0.653 |
| Ramachandran favoured | 99.10 | 99.55 | 99.55 | 99.55 | 99.55 |
| Ramachandran outliers | 0 | 0 | 0 | 0 | 0 |

All datasets are in space group C222 with α = β = γ = 90°.

### Supplementary Table 5

Final refined occupancies used for Fig. 4b and Supplementary Fig. 6.

| **Series** | **Time bin (ms)** | **Retinal–K216 D** | **R82 D** | **w402 A** | **w406 A** |
| --- | --- | --- | --- | --- | --- |
| 75 ms | 7.5–15 | 0.364 | 0.298 | 0.709 | 0.700 |
| 75 ms | 15–22.5 | 0.369 | 0.382 | 0.766 | 0.715 |
| 75 ms | 22.5–30 | 0.321 | 0.339 | 0.864 | 0.792 |
| 75 ms | 30–37.5 | 0.353 | 0.382 | 0.719 | 0.704 |
| 75 ms | 37.5–45 | 0.335 | 0.337 | 0.721 | 0.709 |
| 75 ms | 45–52.5 | 0.290 | 0.343 | 0.791 | 0.742 |
| 75 ms | 52.5–60 | 0.286 | 0.345 | 0.744 | 0.734 |
| 75 ms | 60–67.5 | 0.298 | 0.290 | 0.708 | 0.705 |
| 75 ms | 67.5–75 | 0.259 | 0.330 | 0.853 | 0.759 |
| 500 ms | 0–100 | 0.290 | 0.342 | 0.729 | 0.716 |
| 500 ms | 100–200 | 0.254 | 0.353 | 0.766 | 0.733 |
| 500 ms | 200–300 | 0.249 | 0.358 | 0.790 | 0.741 |
| 500 ms | 300–400 | 0.229 | 0.316 | 0.751 | 0.725 |
| 500 ms | 400–500 | 0.223 | 0.325 | 0.857 | 0.850 |

### Supplementary Table 6

Data collection and refinement statistics for the five 100-ms bins.

| **Segment** | **1** | **2** | **3** | **4** | **5** |
| --- | --- | --- | --- | --- | --- |
| Data collection statistics | | | | | |
| Images taken | 1–10 | 11–20 | 21–30 | 31–40 | 41–50 |
| Time interval, ms | 0–100 | 100–200 | 200–300 | 300–400 | 400–500 |
| Segments merged/collected | 424/790 | 542/790 | 542/790 | 542/790 | 542/790 |
| Total number of images used | 4,240 | 5,420 | 5,420 | 5,420 | 5,420 |
| Space group | C222 | C222 | C222 | C222 | C222 |
| Unit cell |  |  |  |  |  |
| a, Å | 118.41 | 118.15 | 118.21 | 118.60 | 118.41 |
| b, Å | 120.70 | 120.11 | 120.38 | 120.79 | 120.82 |
| c, Å | 36.89 | 36.73 | 36.78 | 36.89 | 36.87 |
| Resolution, Å | 30 – 2.00 (2.05 – 2.00)* | 30 – 2.00 (2.05 – 2.00) | 30 – 2.00 (2.05 – 2.00) | 30 – 2.00 (2.05 – 2.00) | 30 – 2.00 (2.05 – 2.00) |
| Total number of observations/unique | 243,475/18,328 (17,710/1,334) | 308,311/18,142 (23,236/1,331) | 310,084/18,237 (23,292/1,327) | 311,675/18,415 (22,401/1,353) | 311,185/18,387 (22,653/1,350) |
| Completeness, % | 99.6 (99.4) | 99.7 (99.6) | 99.8 (99.9) | 99.8 (99.8) | 99.8 (99.9) |
| Multiplicity | 13 (13) | 17 (17) | 17 (18) | 17 (17) | 17 (17) |
| <(I)/σ(I)> | 8.7 (1.3) | 9.8 (1.4) | 9.6 (1.3) | 9.4 (1.2) | 9.2 (1.1) |
| CC(1/2) | 0.992 (0.552) | 0.998 (0.538) | 0.990 (0.616) | 0.998 (0.511) | 0.993 (0.511) |
| Rmeas | 0.257 (1.817) | 0.203 (1.862) | 0.242 (2.066) | 0.235 (2.026) | 0.260 (2.129) |
| Refinement statistics | | | | | |
| PDB ID | 46OJ | 46OK | 46OL | 46OM | 46ON |
| Resolution, Å | 29.60 – 2.00 | 29.54 – 2.00 | 29.55 – 2.00 | 29.65 – 2.00 | 29.60 – 2.00 |
| Rwork/Rfree | 0.1876/0.2181 | 0.1846/0.2178 | 0.1834/0.2224 | 0.1864/0.2238 | 0.1832/0.2174 |
| No. reflections work/free | 17,428/886 | 17,260/868 | 17,347/876 | 17,515/886 | 17,477/888 |
| No. of atoms |  |  |  |  |  |
| Protein | 3,438 | 3,438 | 3,438 | 3,438 | 3,438 |
| Retinal | 40 | 40 | 40 | 40 | 40 |
| Solvent | 67 | 67 | 67 | 67 | 67 |
| Lipids | 216 | 216 | 216 | 216 | 216 |
| B-factors, Å² |  |  |  |  |  |
| Protein | 32.92 | 32.92 | 32.92 | 32.92 | 32.92 |
| Retinal | 22.98 | 22.98 | 22.98 | 22.98 | 22.98 |
| Solvent | 43.48 | 43.48 | 43.48 | 43.48 | 43.48 |
| Lipids | 62.39 | 62.39 | 62.39 | 62.39 | 62.39 |
| R.m.s. |  |  |  |  |  |
| Bonds, Å | 0.002 | 0.004 | 0.003 | 0.003 | 0.002 |
| Angles, ° | 0.476 | 0.659 | 0.515 | 0.604 | 0.518 |
| Ramachandran favoured | 99.55 | 99.10 | 99.55 | 99.55 | 99.55 |
| Ramachandran outliers | 0 | 0 | 0 | 0 | 0 |

Values in parentheses refer to the highest-resolution shell. All datasets are in space group C222 with α = β = γ = 90°. For the 0–100-ms bin, additional clustering at CC vicinity 0.5 retained 424 of 542 segments, increasing the overall CC₁/₂ from 0.104 to 0.992 (Methods).

### Supplementary Table 7

Data collection and refinement for the five dark dose-control bins.

| **Segment** | **1** | **2** | **3** | **4** | **5** |
| --- | --- | --- | --- | --- | --- |
| Data collection statistics | | | | | |
| Images taken | 1–10 | 11–20 | 21–30 | 31–40 | 41–50 |
| Mean DWD, kGy | 6.5 | 12.0 | 17.5 | 23.0 | 28.5 |
| Segments merged/collected | 309/349 | 309/349 | 309/349 | 308/349 | 309/349 |
| Total number of images used | 3,090 | 3,090 | 3,090 | 3,080 | 3,090 |
| Space group | C222 | C222 | C222 | C222 | C222 |
| Unit cell |  |  |  |  |  |
| a, Å | 117.67 | 117.31 | 117.32 | 117.39 | 117.20 |
| b, Å | 120.42 | 120.28 | 120.21 | 120.25 | 120.17 |
| c, Å | 36.84 | 36.81 | 36.79 | 36.81 | 36.79 |
| Resolution, Å | 30 – 2.00 (2.05 – 2.00)* | 30 – 2.00 (2.05 – 2.00) | 30 – 2.00 (2.05 – 2.00) | 30 – 2.00 (2.05 – 2.00) | 30 – 2.00 (2.05 – 2.00) |
| Total number of observations/unique | 175,837/18,093 (13,220/1,325) | 173,312/18,016 (12,851/1,310) | 174,274/17,984 (12,973/1,300) | 173,475/17,918 (12,673/1,295) | 172,104/17,961 (12,805/1,301) |
| Completeness, % | 99.3 (99.5) | 99.4 (99.8) | 99.3 (99.8) | 98.8 (98.9) | 99.3 (99.5) |
| Multiplicity | 10 (10) | 10 (10) | 10 (10) | 10 (10) | 10 (10) |
| <(I)/σ(I)> | 7.0 (1.1) | 6.8 (1.1) | 7.0 (1.2) | 6.7 (1.1) | 6.4 (0.9) |
| CC(1/2) | 0.976 (0.481) | 0.995 (0.502) | 0.977 (0.457) | 0.989 (0.383) | 0.980 (0.323) |
| Rmeas | 0.260 (2.041) | 0.233 (2.137) | 0.242 (2.066) | 0.242 (2.241) | 0.270 (2.384) |
| Refinement statistics | | | | | |
| PDB ID | 46OO | 46OP | 46OQ | 46OR | 46OS |
| Resolution, Å | 29.42 – 2.00 | 29.33 – 2.00 | 29.33 – 2.00 | 29.35 – 2.00 | 29.30 – 2.00 |
| Rwork/Rfree | 0.1774/0.2157 | 0.1730/0.2090 | 0.1772/0.2153 | 0.1778/0.2172 | 0.1871/0.2296 |
| No. reflections work/free | 17,202/870 | 17,138/862 | 17,101/862 | 17,038/859 | 17073/860 |
| No. of atoms |  |  |  |  |  |
| Protein | 1,873 | 1,873 | 1,873 | 1,873 | 1,873 |
| Retinal | 40 | 40 | 40 | 40 | 40 |
| Solvent | 50 | 50 | 50 | 50 | 50 |
| Lipids | 216 | 216 | 216 | 216 | 216 |
| B-factors, Å² |  |  |  |  |  |
| Protein | 37.49 | 38.86 | 37.93 | 39.68 | 39.42 |
| Retinal | 26.53 | 27.22 | 25.79 | 28.51 | 27.73 |
| Solvent | 49.35 | 50.37 | 49.86 | 51.68 | 50.79 |
| Lipids | 63.01 | 66.85 | 64.07 | 68.93 | 63.31 |
| R.m.s. |  |  |  |  |  |
| Bonds, Å | 0.002 | 0.006 | 0.003 | 0.005 | 0.003 |
| Angles, ° | 0.587 | 0.869 | 0.648 | 0.778 | 0.605 |
| Ramachandran favoured | 99.55 | 99.55 | 99.55 | 99.55 | 99.55 |
| Ramachandran outliers | 0 | 0 | 0 | 0 | 0 |

Values in parentheses refer to the highest-resolution shell. All datasets are in space group C222 with α = β = γ = 90°. Mean diffraction-weighted doses (DWDs) were averaged across each ten-frame subset while retaining prior mesh irradiation and all preceding exposures. They are not dose increments or end-of-subset doses. Values are representative RADDOSE-3D estimates rather than exact crystal-specific doses: measured fluxes varied from 3 × 10¹¹ to 6 × 10¹¹ photons s⁻¹ and crystal dimensions from 20 to 100 µm along each axis (Methods).

### Supplementary Table 8

Selected published serial crystallography datasets for microbial rhodopsins, including time-resolved, continuously illuminated and reference datasets. Counts are diffraction frames, indexed images/patterns or lattices, or merged patterns, as defined in the cited studies.

| **Ref.** | **Protein** | **X-ray source** | **Delivery** | **PDB** | **Time window / condition** | **Merge inputs** | **Data resolution (Å)** |
| --- | --- | --- | --- | --- | --- | --- | --- |
| 8 | BR_WT_ | SACLA BL3 | Jet | 5B6V | Ground | 154,222 | 2.00 |
| 8 | BR_WT_ | SACLA BL3 | Jet | 5B6W | 16 ns | 9,493 | 2.10 |
| 8 | BR_WT_ | SACLA BL3 | Jet | 5B6X | 760 ns | 10,054 | 2.10 |
| 8 | BR_WT_ | SACLA BL3 | Jet | 5B6Y | 36.2 µs | 10,011 | 2.10 |
| 8 | BR_WT_ | SACLA BL3 | Jet | 5B6Z | 1.725 ms | 8,375 | 2.10 |
| 8 | BR_WT_ | SACLA BL3 | Jet | 5H2H | 40 ns | 9,448 | 2.10 |
| 8 | BR_WT_ | SACLA BL3 | Jet | 5H2I | 110 ns | 10,947 | 2.10 |
| 8 | BR_WT_ | SACLA BL3 | Jet | 5H2J | 290 ns | 8,578 | 2.10 |
| 8 | BR_WT_ | SACLA BL3 | Jet | 5H2K | 2 µs | 11,215 | 2.10 |
| 8 | BR_WT_ | SACLA BL3 | Jet | 5H2L | 5.25 µs | 9,028 | 2.10 |
| 8 | BR_WT_ | SACLA BL3 | Jet | 5H2M | 13.8 µs | 9,254 | 2.10 |
| 8 | BR_WT_ | SACLA BL3 | Jet | 5H2N | 95.2 µs | 11,009 | 2.10 |
| 8 | BR_WT_ | SACLA BL3 | Jet | 5H2O | 250 µs | 10,578 | 2.10 |
| 8 | BR_WT_ | SACLA BL3 | Jet | 5H2P | 657 µs | 9,596 | 2.10 |
| 9 | BR_WT_ | LCLS CXI | Jet | 6GA1 | Ground | 22,485 | 1.70 |
| 9 | BR_WT_ | LCLS CXI | Jet | 6GA2 | Ground | 69,518 | 1.80 |
| 9 | BR_WT_ | LCLS CXI | Jet | 6GA3 | 33 ms | 9,022 | 2.10 |
| 9 | BR_WT_ | LCLS CXI | Jet | 6GA4 | 1 ps | 11,590 | 1.80 |
| 9 | BR_WT_ | LCLS CXI | Jet | 6GA5 | 3 ps | 9,739 | 1.90 |
| 9 | BR_WT_ | LCLS CXI | Jet | 6GA6 | 10 ps | 8,997 | 1.80 |
| 9 | BR_WT_ | LCLS CXI | Jet | 6GA7 | 0.24 ps | 10,000 | 1.80 |
| 9 | BR_WT_ | LCLS CXI | Jet | 6GA8 | 0.33 ps | 10,000 | 1.80 |
| 9 | BR_WT_ | LCLS CXI | Jet | 6GA9 | 0.39 ps | 10,000 | 1.80 |
| 9 | BR_WT_ | LCLS CXI | Jet | 6GAA | 0.43 ps | 10,000 | 1.80 |
| 9 | BR_WT_ | LCLS CXI | Jet | 6GAB | 0.46 ps | 10,000 | 1.80 |
| 9 | BR_WT_ | LCLS CXI | Jet | 6GAC | 0.49 ps | 10,000 | 1.80 |
| 9 | BR_WT_ | LCLS CXI | Jet | 6GAD | 0.53 ps | 10,000 | 1.80 |
| 9 | BR_WT_ | LCLS CXI | Jet | 6GAE | 0.56 ps | 10,000 | 1.80 |
| 9 | BR_WT_ | LCLS CXI | Jet | 6GAF | 0.59 ps | 10,000 | 1.80 |
| 9 | BR_WT_ | LCLS CXI | Jet | 6GAG | 0.63 ps | 10,000 | 1.80 |
| 9 | BR_WT_ | LCLS CXI | Jet | 6GAH | 0.68 ps | 10,000 | 1.80 |
| 9 | BR_WT_ | LCLS CXI | Jet | 6GAI | 0.74 ps | 10,000 | 1.80 |
| 9 | BR_WT_ | LCLS CXI | Jet | 6RMK | Ground (re-refined) | 10,000 | 1.80 |
| 10 | BR_WT_ | LCLS CXI | Jet | 6G7H | Ground | 241,475 | 1.50 |
| 10 | BR_WT_ | LCLS CXI | Jet | 6G7I | 49-406 fs | 54,479 | 1.50 |
| 10 | BR_WT_ | LCLS CXI | Jet | 6G7J | 457-646 fs | 54,955 | 1.50 |
| 10 | BR_WT_ | LCLS CXI | Jet | 6G7K | 10 ps | 30,245 | 1.50 |
| 10 | BR_WT_ | LCLS CXI | Jet | 6G7L | 8.33 ms | 235,238 | 1.55 |
| 11 | BR_WT_ | SLS PXI | Jet | 6RNJ | 0-5 ms | 13,346* | 2.30 |
| 11 | BR_WT_ | SLS PXI | Jet | 6RPH | 10-15 ms | 13,384* | 2.30 |
| 11 | BR_WT_ | SLS PXI | Jet | 6RQO | Continuous | 114,052 | 1.80 |
| 11 | BR_WT_ | SLS PXI | Jet | 6RQP | Ground | 119,374 | 1.80 |
| 12 | BR_WT_ | SACLA BL3 | Jet | 9F9B | Ground | 325,179 | 1.83 |
| 12 | BR_WT_ | SACLA BL3 | Jet | 9F9C | 16 ns | 25,487 | 1.95 |
| 12 | BR_WT_ | SwissFEL ESA | Jet | 9F9D | Ground | 32,765 | 1.65 |
| 12 | BR_WT_ | SwissFEL ESA | Jet | 9F9E | 6 ps | 43,892 | 1.71 |
| 12 | BR_WT_ | SwissFEL ESA | Jet | 9F9F | 6 ps | 45,778 | 1.66 |
| 12 | BR_WT_ | LCLS CXI | Jet | 9F9G | Ground | 241,475 | 1.44 |
| 12 | BR_WT_ | LCLS CXI | Jet | 9F9H | 10 ps | 30,245 | 1.55 |
| 12 | BR_WT_ | SACLA BL3 | Jet | 9F9I | Ground | 178,536 | 1.59 |
| 12 | BR_WT_ | SACLA BL3 | Jet | 9F9J | 10 ps | 32,400 | 1.65 |
| 13 | Arch-3 | DIAMOND I24 | Jet | 6GUY | Ground | 1,438 | 2.20 |
| 13 | Arch-3 | SACLA BL2 | Jet | 7ZY3 | 110 ns | 11,912 | 1.70 |
| 14 | BcXeR | DESY P14 | Jet | 7ZNE | Ground | 67,912 | 2.10 |
| 14 | BcXeR | DESY P14 | Jet | 7ZNI | 7.5-15 ms | 12,449 | 2.20 |
| 14 | BcXeR | DESY P14 | Jet | 7ZNG | Ground | 17,997 | 2.30 |
| 14 | BcXeR | DESY P14 | Jet | 7ZNH | 250-750 µs | 17,526 | 2.30 |
| 15 | KR2 | ALBA XALOC | Jet | 6YC0 | Continuous | 38,761 | 2.50 |
| 16 | KR2 | SwissFEL ESA | Jet | 6TK1 | 20 ms | 48,329 | 2.10×2.20×1.60 |
| 16 | KR2 | SwissFEL ESA | Jet | 6TK2 | 1 ms | 60,629 | 2.10×2.20×1.70 |
| 16 | KR2 | SwissFEL ESA | Jet | 6TK3 | 30 + 150 µs | 50,754 + 48,383 | 2.00×2.20×1.60 |
| 16 | KR2 | SwissFEL ESA | Jet | 6TK4 | 1 + 16 ns | 45,045 + 50,165 | 2.00×2.20×1.60 |
| 16 | KR2 | SwissFEL ESA | Jet | 6TK5 | 800 fs + 2 ps | 47,801 + 47,595 | 2.00×2.20×1.60 |
| 16 | KR2 | SwissFEL ESA | Jet | 6TK6 | Ground (neutral pH) | 158,832 | 2.20×2.30×1.60 |
| 16 | KR2 | SwissFEL ESA | Jet | 6TK7 | Ground (acidic pH) | 47,518 | 2.00×2.30×1.60 |
| 17 | KR2 | SwissFEL ESA | Jet | 8CL7 | Ground | 53,745 | 1.98 |
| 17 | KR2 | SwissFEL ESA | Jet | 8CL8 | 1 µs | 25,036 | 2.04 |
| 18 | ClR | LCLS CXI | Jet | 7CRI | 1 ps | 29,469 | 1.73 |
| 18 | ClR | LCLS CXI | Jet | 7CRJ | Ground | 37,044 | 1.65 |
| 18 | ClR | LCLS CXI | Jet | 7CRK | 2 ps | 25,690 | 1.72 |
| 18 | ClR | LCLS CXI | Jet | 7CRL | 50 ps | 23,718 | 1.77 |
| 18 | ClR | LCLS CXI | Jet | 7CRS | 100 ps | 35,106 | 1.74 |
| 18 | ClR | LCLS CXI | Jet | 7CRT | 100 ps | 47,795 | 1.74 |
| 18 | ClR | LCLS CXI | Jet | 7CRX | 100 ps | 22,474 | 1.77 |
| 18 | ClR | LCLS CXI | Jet | 7CRY | 100 ps | 41,253 | 1.81 |
| 19 | NM-R3 | SACLA BL3 | Jet | 7VGT | Ground | 31,017 | 2.10 |
| 19 | NM-R3 | SACLA BL3 | Jet | 7VGU | 1 ms | 46,447 | 2.10 |
| 19 | NM-R3 | SACLA BL3 | Jet | 7VGV | Ground (anion-depleted) | 25,936 | 2.30 |
| 20 | NmHR | SwissFEL ESA | Jet | 7O8F | Ground | 228,156 | 1.45 |
| 20 | NmHR | SwissFEL ESA | Jet | 7O8G | 10 ps | 116,249 | 1.45 |
| 20 | NmHR | SwissFEL ESA | Jet | 7O8H | 10 ns | 107,514 | 1.45 |
| 20 | NmHR | SwissFEL ESA | Jet | 7O8I | 1 µs | 112,288 | 1.45 |
| 20 | NmHR | SwissFEL ESA | Jet | 7O8J | 20 µs | 113,241 | 1.45 |
| 20 | NmHR | SwissFEL ESA | Jet | 7O8K | 300 µs | 102,394 | 1.45 |
| 20 | NmHR | SLS PXI | Jet | 7O8L | Ground reference (80–200 ms) | 2,299,838* | 1.80 |
| 20 | NmHR | SLS PXI | Jet | 7O8M | 0-5 ms | 94,362* | 2.00 |
| 20 | NmHR | SLS PXI | Jet | 7O8N | 5-10 ms | 92,645* | 2.00 |
| 20 | NmHR | SLS PXI | Jet | 7O8O | 10-15 ms | 92,555* | 2.00 |
| 20 | NmHR | SLS PXI | Jet | 7O8P | 15-20 ms | 94,666* | 2.00 |
| 20 | NmHR | SLS PXI | Jet | 7O8Q | 20-25 ms | 95,514* | 2.00 |
| 20 | NmHR | SLS PXI | Jet | 7O8R | 25-30 ms | 94,587* | 2.00 |
| 20 | NmHR | SLS PXI | Jet | 7O8S | 30-35 ms | 95,381* | 2.00 |
| 20 | NmHR | SLS PXI | Jet | 7O8T | 35-40 ms | 93,890* | 2.00 |
| 20 | NmHR | SLS PXI | Jet | 7O8U | 40-50 ms | 185,285* | 2.00 |
| 20 | NmHR | SLS PXI | Jet | 7O8V | 50-60 ms | 188,656* | 2.00 |
| 20 | NmHR | SLS PXI | Jet | 7O8Y | Ground | 300,879 | 1.75 |
| 20 | NmHR | SLS PXI | Jet | 7O8Z | Continuous | 734,749 | 1.80 |
| 21 | C1C2 | SACLA BL3 | Jet | 7C86 | Ground | 62,218 | 2.30 |
| 21 | C1C2 | SACLA BL3 | Jet | 7E6Y | 1 µs | 16,924 | 2.50 |
| 21 | C1C2 | SACLA BL3 | Jet | 7E6Z | 50 µs | 21,129 | 2.50 |
| 21 | C1C2 | SACLA BL3 | Jet | 7E70 | 250 µs | 16,442 | 2.50 |
| 21 | C1C2 | SACLA BL3 | Jet | 7E71 | 1 ms | 22,156 | 2.50 |
| 21 | C1C2 | SACLA BL3 | Jet | 7E6X | 4 ms | 24,015 | 2.50 |
| 22 | C1C2 | SLS PXI | Jet | 9GO1 | Ground | 55,249 | 2.60 |
| 22 | C1C2 | SLS PXI | Jet | 9GO2 | Continuous illumination (~100 ms) | 43,646 | 2.70 |
| 23 | NpSRII | SLS PXI | Jet | 8PWG | 90-120 ms | 7,163* | 2.40 |
| 23 | NpSRII | SLS PXI | Jet | 8PWI | 60-90 ms | 6,890* | 2.40 |
| 23 | NpSRII | SLS PXI | Jet | 8PWJ | 30-60 ms | 6,577* | 2.40 |
| 23 | NpSRII | SLS PXI | Jet | 8PWP | 0-30 ms | 6,481* | 2.40 |
| 23 | NpSRII | SLS PXI | Jet | 8PWQ | 120-150 ms | 7,705* | 2.40 |
| 23 | NpSRII | SLS PXI | Jet | 9H1W | Grouped dark reference (150–240 ms) | 24,485* | 2.40 |
| 23 | NpSRII | SLS PXI | Jet | 9H1X | Continuous | 21,518 | 1.85 |
| 23 | NpSRII | SLS PXI | Jet | 9H20 | Ground | 26,382 | 2.20 |

Resolution values are diffraction-data limits and may differ from model-refinement limits. Anisotropic limits are reproduced as reported. Multiple time windows or multiple datasets may share contributing crystals. These entries are contextual and are not a matched comparison with BR_MT_. Arch-3, archaerhodopsin-3; BcXeR, Bacillus coahuilensis xenorhodopsin; KR2, Krokinobacter eikastus rhodopsin 2; ClR and NmHR/NM-R3, chloride-pumping rhodopsins; C1C2, chimeric channelrhodopsin; NpSRII, Natronomonas pharaonis sensory rhodopsin II. The ultrafast BR_WT_ entries attributed to Nass Kovacs *et al*. include overlapping temporal bins; their counts must not be summed as independent crystal populations. Datasets 9F9B/C and 9F9G/H reprocess the Nango *et al*. and Nogly *et al*. data, respectively, and do not represent independent acquisitions. An asterisk marks repeated temporal measurements from a crystal. Asterisks do not imply that the same crystals contribute to every time window.

### Supplementary Table 9

Refinement statistics for the K-like and M-like states refined against extrapolated amplitudes.

| **Time window, ms** | **0–7.5** | **7.5–15** |
| --- | --- | --- |
| Intermediate | K-like | M-like |
| PDB ID | 46NX | 46NY |
| Resolution, Å | 11.91 – 2.00 | 11.96 – 2.01 |
| Rwork/Rfree | 0.3849/0.3996 | 0.2826/0.3157 |
| No. reflections work/free | 16,488/869 | 16,560/870 |
| No. of atoms |  |  |
| Protein | 1,721 | 1,717 |
| Retinal | 20 | 20 |
| Solvent | 40 | 26 |
| Lipids | 0 | 216 |
| B–factors, Å² |  |  |
| Protein | 38.97 | 47.56 |
| Retinal | 31.51 | 38.41 |
| Solvent | 43.72 | 52.38 |
| Lipids | — | 66.38 |
| R.m.s. |  |  |
| Bonds, Å | 0.002 | 0.002 |
| Angles, ° | 0.469 | 0.456 |
| Ramachandran favoured | 99.55 | 99.10 |
| Ramachandran outliers | 0 | 0 |

K-like and M-like models were refined against qFextr amplitudes at extrapolation fractions of 0.20 and 0.30, respectively. These correspond to extrapolation factors α = 1/f of 5.0 and 3.33. The M-like fraction maximized the integrated difference-density signal. The K-like fraction coincided with the maximum Pearson map correlation, whereas difference-map maximization selected 0.10; the criteria and scans are described in Methods.
